# Low-cost rhizotron imaging and zero-shot deep-learning resolve temporal, spatial, and genetic variation in grapevine rootstock root systems

**DOI:** 10.64898/2026.09.10.750709

**Authors:** Jose R. Munoz, Efrain Torres-Lomas, Sadikshya Sharma, Yaniv Lupo, Andrew J. McElrone, Luis Diaz-Garcia

## Abstract

Root system architecture shapes how grapevine rootstocks take up water and nutrients, yet roots remain the least phenotyped grapevine organ because they are hidden and hard to image. We present a low-cost phenotyping pipeline that pairs custom acrylic rhizotrons (about US$30 each) with a consumer flatbed scanner and BiRefNet, a general-purpose deep-learning model used without training on root images, followed by automated mask cleaning, skeleton-based trait extraction, and soil moisture mapping. We tested it on nine commercial rootstocks scanned 16 times over 42 days after transplanting (DAT), with half under a ten-day water deficit. From 1,108 images we extracted 21 whole-root, depth-resolved, and topological traits. Genotypes differed in nearly every trait and in how they changed over time. Heritability of size and branching traits peaked at 0.92-0.93 between 21 and 31 DAT and fell for width, depth, and convex hull once roots reached the rhizotron walls, defining the best measurement window. The image-derived soil moisture map accurately tracked the deficit and its recovery. Deficit plants shifted new root growth to deeper soil without growing less overall, and the substrate dried fastest around older and denser roots. Root brightness decreased with root age and local moisture, and transport segments (axes serving several tips) were brighter than terminal laterals in every genotype. Root system size was associated with stomatal conductance in well-watered plants, and stomatal recovery after re-watering correlated with new root growth. The pipeline turns simple hardware into a quantitative, time-resolved root phenotyping platform suitable for breeding.

## Introduction

Water availability is a major limiting factor in viticulture, particularly in regions with a Mediterranean climate where irregular rainfall often leads to prolonged drought [1,2]. Using drought-tolerant grapevine rootstocks is crucial in mitigating the adverse effects of water limitation on grapevine growth and productivity [3–5]. Commercial rootstocks vary substantially in their drought tolerance [6,7], a trait closely associated with root system architecture (RSA). RSA, defined as the spatial arrangement of primary, lateral, and fine roots within the soil, influences the grapevine’s water and nutrient uptake, anchorage, and overall stress resilience [8–10]. RSA therefore plays a crucial role in enabling grapevines to maintain productivity under drought, making it a key target for rootstock breeding programs [5,11,12]. Studies comparing drought-tolerant and susceptible rootstocks have shown that drought-tolerant rootstocks typically have traits that support water uptake from deeper soil layers with more stable water reserves [13–17]. For instance, drought-tolerant rootstocks often have steeper root angles [4], thicker fine roots [18], and plastic root growth responses to spatial and temporal variation in soil moisture [14]. These characteristics direct root growth into deeper, wetter soil and support the rapid growth of new roots in rewetted soil, enabling plants to quickly resume water uptake after a drought [19,20]. Although RSA is essential for drought adaptation, characterizing root traits remains challenging because roots are hidden and existing phenotyping methods have practical limits.

Root systems remain the most challenging plant organs to phenotype, especially for perennials under field conditions, because of their subterranean nature and long juvenile periods [21]. Many methods, both destructive and non-destructive, have been developed to study their structure and function, but most are expensive and labor-intensive, which makes them poorly suited to large breeding populations. Traditional techniques such as soil coring [22], trenching [23], and “shovelomics” [24] are destructive and time-consuming, typically involving the excavation and washing of root systems, during which fine roots are lost. Non-destructive alternatives include nuclear magnetic resonance imaging [25], X-ray computed tomography [26–28], neutron radiography [19], and rhizotrons or rhizoboxes [9,29–32]. Magnetic resonance imaging, neutron radiography, and computed tomography are non-invasive, but their cost and slow data collection make them impractical for breeding programs. Rhizotrons, in contrast, are affordable and allow repeated imaging of the same root system with cameras or flatbed scanners, capturing growth dynamics over time and scaling to larger populations.

While rhizotrons provide a simple and accessible way to image roots repeatedly, extracting quantitative traits from the images remains the bottleneck. Several software tools are available for root image analysis, including WinRHIZO [33], RootNav [34], SmartRoot [35], and RhizoVision Explorer [36], among others. However, many of these programs are constrained by cost, by semi-automated workflows that require substantial manual input or annotation, or by reduced performance on complex root systems growing in dark, heterogeneous media [21]. Recent advances in computer vision, particularly deep-learning models for high-resolution image segmentation, offer a way past these limitations. The bilateral reference network (BiRefNet) [37] was designed to separate thin, high-resolution objects from their background and was trained on large, general-purpose image datasets. Because it does not require task-specific training, BiRefNet can, in principle, segment complex root systems fully automatically, with no manual annotation at all.

Here we describe a cost-effective root phenotyping pipeline that combines simple, custom-made rhizotrons and a flatbed scanner with zero-shot BiRefNet segmentation, automated mask cleaning, and skeleton-graph trait extraction. To evaluate the pipeline, we grew nine commercial grapevine rootstocks in rhizotrons for six weeks, imaged them three times per week, and imposed a water deficit on half of the plants. We asked whether image-derived traits (i) resolve genotype differences and their temporal dynamics, (ii) capture the vertical deployment of roots and its plasticity under water deficit, (iii) relate to soil moisture and to shoot physiology, and (iv) carry enough genetic signal, and early enough, to be useful for selection in breeding populations.

## Materials and Methods

### Experimental and technical design

The study combines a greenhouse experiment with an image-analysis pipeline, both summarized in Figure 1. Nine commercial grapevine rootstocks were rooted from dormant cuttings and grown in 110 custom acrylic rhizotrons; 80 plants produced usable images and form the analysis set. Each rhizotron was scanned on a flatbed scanner three times per week for six weeks (16 imaging dates ranging from 6 to 42 days after transplanting), and half of the plants were subjected to a ten-day water deficit starting at 29 DAT. The technical pipeline proceeds in five steps: scans of the same rhizotron are aligned to a common frame; roots are segmented with BiRefNet without any task-specific training; masks are cleaned with automated rules; traits are extracted from the cleaned masks and their skeleton graphs; and the substrate visible in the same scans is converted into a soil moisture map. All traits, statistics, and figures were computed from the cleaned masks with the same documented code. The sections below describe each component in detail.

**Figure 1.**
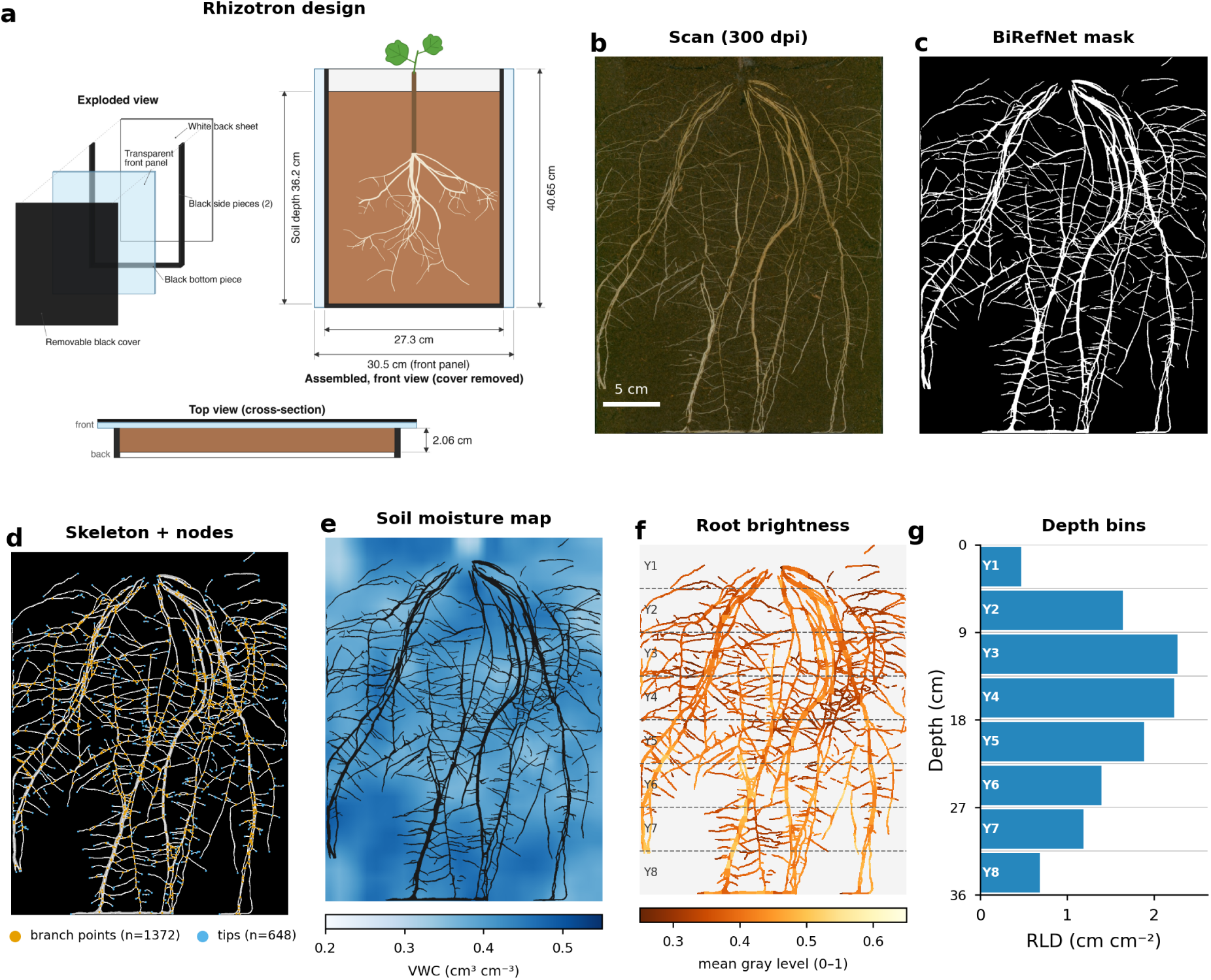
Rhizotron design and image-analysis pipeline. (a) Exploded and assembled views of the acrylic rhizotron with component dimensions. (b) Flatbed scan (300 dpi) of the front panel of a Freedom rootstock rhizotron at 40 DAT (water-deficit treatment); scale bar 5 cm. (c) Zero-shot BiRefNet mask of the same scan (threshold 0.3). (d) Skeleton of the cleaned mask (white) with branch points (orange) and root tips (blue). (e) Image-derived soil moisture map (volumetric water content estimated from substrate gray level in 0.9 × 0.85 cm cells, smoothed) with roots overlaid. (f) Root pixel brightness (mean gray level of eroded root pixels per cell) painted on the mask, with the eight depth bins (Y1-Y8) used for spatial analyses. (g) Root length density in the same bins.

### Plant material and experimental set-up

The pipeline was evaluated using commercially available grapevine rootstocks in a greenhouse trial conducted in spring 2025 at the University of California, Davis. Dormant cuttings were collected in February 2025 from mother vines located at the UC Davis Tyree research vineyard (Davis, California). The rootstocks were 110R and 1103P (*V. berlandieri* × *V. rupestris*), 101-14 Mgt and 3309C (*V. riparia* × *V. rupestris*), 5C (*V. berlandieri* × *V. riparia*), Ramsey (*V. champinii*), Freedom (1613C × Dog Ridge), GRN-1 (*V. rupestris* × *Muscadinia rotundifolia*) and GRN-3 ((*V. rufotomentosa* × (Dog Ridge × Riparia Gloire)) × *V. champinii* c9038). Two additional rootstocks (420A and O39-16) were included at planting but failed to root from dormant cuttings and were excluded from all analyses. Cuttings were soaked overnight in a 1% bleach solution, placed in callusing medium, and kept in a dark room at 100% relative humidity and 29 °C for approximately two weeks. Once sufficient callus developed, the top buds were waxed, and the cuttings were transplanted into rhizotrons filled with a 1:1 (v/v) coco coir:peat moss mixture. Transplanted cuttings were returned to the dark room until bud break, then moved to a fog room (100% relative humidity, 27 °C) for approximately four days, and finally transferred to a greenhouse, where the rhizotrons were placed in racks at approximately 45° with the transparent panel face down, so that roots grew along the front panel. The front panel was kept covered with the removable black cover except during scanning, which excludes light from the root zone and limits algal growth. Plants remained under shade netting for about one week to acclimate. Rhizotrons were arranged in a completely randomized design with up to ten replicates per genotype. Transplanting is considered day 0 (14 March 2025); the first scan was collected 6 days after transplanting (DAT).

### Rhizotron construction and imaging

Rhizotrons were assembled from cut-to-size cast acrylic (TAP Plastics, Stockton, CA, USA) bonded with acrylic cement (Figure 1a). Each rhizotron consists of two black side pieces and a black bottom piece, a white back sheet that limits solar heating of the substrate, and a transparent front panel (30.5 × 40.65 cm) that extends beyond the sides so that a removable black cover can be attached to exclude light from the root zone. The internal cavity is 27.3 cm wide and 2.06 cm thick and holds a substrate column about 36.2 cm deep; the scanned area analyzed in this study covers 27.3 × 35.8 cm of substrate. Material cost was approximately US$30 per rhizotron.

Roots were imaged through the front panel with an OpticPro A320L A3 flatbed scanner (Plustek, Santa Fe Springs, CA, USA) at 300 dpi (0.085 mm per pixel), which takes approximately 10 s per rhizotron; the same scanner can resolve root hairs at 1,600 dpi (Figure S1). An ArUco fiducial marker fixed to each front panel identified the rhizotron and provided a scale reference. Scans were collected three times per week (Monday, Wednesday, Friday) from 20 March to 25 April 2025, giving 16 imaging dates between 6 and 42 DAT and 1,689 scans in total. Condensation on the front panel affected many of the first scans (6 DAT), and these images were used only when the root system was clearly visible.

### Water-deficit treatment

A water-deficit treatment was initiated at 29 DAT and ended by re-watering at 39 DAT. Because cuttings root asynchronously, plants were assigned to the deficit treatment only after they had developed a moderately sized root system, so that the deficit was applied to established root systems; the remaining plants of each genotype were kept well-watered. As a consequence, plants in the deficit group had, on average, larger root systems at the start of the treatment than the well-watered controls, and treatment comparisons were therefore made on within-plant changes and on size-adjusted growth rates rather than on absolute values. Of the 85 plants assigned to a treatment, 80 produced usable root images and form the analysis set: 39 were subjected to the deficit and 41 remained well-watered (7-10 per genotype, with 3-6 per genotype and treatment; Table S1). The assignment to the deficit treatment was made by eye from the scans available at 28 DAT, and no attempt was made to balance root size between groups.

### Physiological measurements

Stomatal conductance to water vapor (gsw) was measured at midday (11:00-13:00) with an LI-600 porometer/fluorometer (LI-COR, Lincoln, NE, USA) on one fully expanded leaf per plant at three time points during the deficit (32, 35 and 38 DAT) and once after re-watering (42 DAT). In total, 318 measurements were obtained from the 80 plants in the root analysis set, 313 of which could be paired with a root image scanned within one day.

### Image alignment and segmentation

Scans of the same rhizotron were aligned to a common frame by translation, using the ArUco marker and phase correlation, so that pixels corresponded across dates. Aligned scans were then segmented with BiRefNet [37], a model for dichotomous (foreground-background) image segmentation. BiRefNet is designed to preserve fine object boundaries and structures, including long, thin features such as hair, branches, and wires, making it well suited to the geometry of roots visible on rhizotron panels. Publicly released general-purpose weights were used without fine-tuning on root images (zero-shot inference; https://huggingface.co/ZhengPeng7/BiRefNet). The model produced a foreground probability map for each image, which was resized to the original scan dimensions and binarized using a threshold of 0.3. A threshold of 0.7 was also evaluated but removed many fine roots and was therefore discarded following visual comparison. Inference was performed on an AWS EC2 G5.2xlarge instance at approximately 1.5 s per image. Each mask (4,230 × 3,346 pixels, including a 60-pixel border added during alignment) was stored as an 8-bit PNG.

### Mask cleaning and quality control

Masks were cleaned automatically in Python 3.11 (scikit-image 0.26 [38], OpenCV 4.13 [39], SciPy 1.17). Connected components smaller than 80 pixels (0.6 mm^2^) were removed as noise. Remaining objects were grouped into clusters by single-linkage at a distance of 6 cm, and clusters that were not the largest were retained only if they started in the upper 35% of the rhizotron (the zone in which adventitious roots emerge from the cutting) or if they contained at least 3% of the pixels of the largest cluster and began within 8 cm of its lower edge; clusters confined to the lowest 30% of the image and disconnected from the main root system (marks left by earlier experiments in four rhizotrons) were removed. This procedure altered a median of 0.13% of mask pixels per image and more than 1% in only 22 of the 1,108 analyzed images.

Images were excluded from analysis when (i) they were listed in a manually curated exclusion file compiled by inspecting every scan (147 images, mostly early dates with condensation or no visible roots), (ii) the cleaned mask contained fewer than 0.1 cm^2^ of roots (28 images), or (iii) root area decreased by more than 25% relative to the previous retained image of the same plant, which indicates a segmentation failure (2 images; the earlier image was flagged instead when the following image was consistent with the trajectory). Of the 1,285 scans of treated plants, 1,108 (86%) were retained, with a mean of 13.9 images per plant (Figure S2).

### Trait extraction

All traits were computed from the cleaned masks. Root area, perimeter (from external contours), convex hull area, and the bounding-box width and depth were measured at full resolution; aspect ratio was width/depth, and solidity was root area divided by convex hull area. Skeleton-based traits were computed at half-resolution (0.17 mm per pixel) after skeletonizing the mask and converting the skeleton into a graph using the skan library [40]. Total root length was the sum of skeleton pixel lengths with diagonal steps weighted by √2. Root tips were skeleton nodes of degree one and branch points were nodes of degree three or more; branching density was branch points per centimeter of root length. For every skeleton branch of at least 0.15 cm, the orientation of the chord joining its two ends was expressed as the angle to the vertical (0° vertical, 90° horizontal), and the length-weighted mean of this angle (mean angle to gravity) and the length fraction with an angle below 30° (vertical fraction) were recorded. Tortuosity was the length-weighted mean ratio of branch length to chord length. Mean root diameter was calculated from root area divided by total root length (the same ratio can be computed for any part of the mask, for example per depth bin). Lateral spread was the standard deviation of root pixel x-coordinates. The rhizotron image was divided into eight equal depth bins of 4.5 cm (Y1, 0-4.5 cm, to Y8, 31.3-35.8 cm) and root length density (RLD, cm of root per cm^2^ of bin area) and root pixel density (fraction of bin area occupied by roots) were calculated per bin; total RLD was total length divided by the imaged soil area. The depth above which 50% of the root length was located (D50) and the depth of the root centroid summarized vertical deployment.

### Segment support (root “traffic”)

Binary masks quantify root area, length, and depth distribution but not root topology, the branching pattern and connectivity of the root system. Annotating root orders (primary, secondary, higher) by hand does not scale and becomes ambiguous at later imaging dates, when laterals proliferate, roots cross and overlap in projection, and masks have gaps. We therefore described topology with a simple, scalable construction on the skeleton rather than on the mask (Supplementary Note S1; Figure S3). Each skeleton was cut at branch points into segments whose ends are tips (free ends of the skeleton, including the end at the cutting and ends created by gaps) or branch points; no hidden connections were inferred. From the tip of every terminal segment (one tip) a path was traced inward through segments bounded by two branch points, selecting at each branch point, among the segments not yet on the path and not leading toward the bottom of the rhizotron, the one with the smallest turning angle, and ending when no admissible segment remained. In the transport-network analogy that gives the metric its name, each path is a route from a root tip toward the cutting, and the support of a segment is its traffic, the number of routes that pass through it: 1 on terminal segments, the number of tips served on shared axes, and 0 on segments that no route selected, whether isolated (two tips, where no route starts) or connected, such as the bridge artifacts created where two roots cross. Support was computed for every image with a non-empty skeleton and summarized for the 1,108 retained images as the length-weighted mean support, the fraction of root length with support ≥ 2 (hereafter transport-root fraction), and the length-weighted distribution of segment angles.

### Image-derived soil moisture and root brightness

Soil moisture was estimated from the gray level of the substrate in the scans converted to grayscale (0 = black, 1 = white). A calibration rhizotron filled with the same substrate was saturated and scanned repeatedly during dry-down while volumetric water content (VWC) was recorded with an ECH2O EC-5 sensor (METER Group, Pullman, WA, USA); a linear model related mean soil gray level (0-1) to VWC (VWC = 1.308 - 4.789 × gray; R^2^ = 0.94, n = 15). For each aligned scan, root pixels and a 15-pixel margin around them were masked out, the soil gray level was averaged within a 40 × 32 grid of cells of about 0.9 × 0.85 cm (these grid cells are the “cells” of all cell-level analyses below) and converted to VWC, and cell values were smoothed with a 3 × 3 moving average to produce a soil moisture map (Figure 1e). The mean VWC of the rhizotron was used as a plant-level covariate. Root brightness was the mean gray level of root pixels in the grayscale image (higher values mean lighter roots), after a one-pixel erosion of the mask to exclude edge pixels, computed per image and per grid cell (cells with at least 100 root pixels). Because scans were aligned across dates, the age of the roots in each cell was estimated as the number of days since roots were first detected in that cell (cells with at least 30 root pixels). Segment brightness was obtained by rasterizing each skeleton segment as a 7-pixel-wide line, intersecting it with the eroded mask and averaging the gray level of the resulting pixels (segments with at least 30 pixels; 426,000 segments from the 80 analyzed plants); the age and local VWC of a segment were taken from the grid cell containing its midpoint.

## Data analysis

Statistical analyses were performed in Python using statsmodels 0.14 [41] (linear mixed models with random intercepts for plant and, where stated, for plant × date). For each of 21 whole-root and topological traits, a mixed model with genotype, imaging day (as a factor), their interaction, and treatment as fixed effects, and plant as a random intercept was fitted by maximum likelihood; size-related traits were log-transformed. Genotype, imaging day, genotype × day, and treatment effects were tested with likelihood-ratio tests against reduced models, and P values were adjusted for the false discovery rate (FDR) across traits. Genotype effects at individual imaging days were tested using ANOVA with treatment as a covariate. Principal component analysis was run on 15 scaled traits (size traits were log-transformed) across all 1,108 plant-date observations. Depth-resolved RLD and root density were analyzed using a mixed model that included genotype, depth bin, their interaction, imaging day, and treatment, with plant as a random effect.

Water-deficit effects were assessed in two ways that account for the larger initial size of the root systems of deficit plants. First, the relative growth rate of root area (RGR, day^-1^) between consecutive images was modeled with treatment, phase (pre-treatment, 6-28 DAT; deficit, 29-38 DAT; recovery, 39-42 DAT), their interaction, genotype, and the log of root area at the start of each interval as fixed effects and plant as a random effect. Second, for each plant imaged at both 28 and 38 DAT (n = 76), the change in each trait over the deficit period was regressed on treatment, genotype, the trait value at 28 DAT (for depth-bin RLD, the bin’s own RLD at 28 DAT) and log total root length at 28 DAT; genotype × treatment interactions were tested by comparing nested models. As a check against residual confounding by size, the analysis was repeated on the 53 plants whose total root length at 28 DAT fell within the range shared by both treatments (152-622 cm; 20 well-watered, 33 deficit). Stomatal conductance was analyzed with a mixed model with treatment, date, their interaction, and genotype as fixed effects and plant as a random effect. Associations between root traits and gsw were tested by pairing each gsw measurement with the root image closest in time (31, 35, 38 and 42 DAT), standardizing each root trait, and fitting mixed models with the standardized trait, treatment, genotype and date as fixed effects and plant as a random effect; P values were FDR-adjusted across the 22 variables tested. The same models were fitted within each treatment group and with log root area as an additional covariate, and the change in gsw between 38 and 42 DAT was regressed on the change in log total root length over the same interval, genotype, and treatment. The relationship between root brightness, local soil moisture, and root age was analyzed at the grid-cell level (a random sample of 40,000 of 271,000 rooted cells) with a mixed model including local VWC, root age, cell depth, treatment and genotype as fixed effects and plant as a random effect; the model was refitted with an additional plant × date random intercept and with soil moisture estimated after excluding a margin of 1.3, 5 or 10 mm around the roots, to assess the contribution of substrate immediately adjacent to the root, and at the level of plant-date means.

Observations at the level of grid cells and skeleton segments are numerous but nested within 80 plants and 16 dates; the P values of these models are therefore reported descriptively, and effect sizes with their standard errors are the quantities of interest. Root-soil moisture coupling was analyzed on all grid cells with sufficient visible substrate (1.30 million cell-date observations from the 1,108 retained images, excluding the outermost columns). Three quantities were derived. The local moisture anomaly of a cell was its VWC minus the mean VWC of all cells in the same row (depth) of the same image; it is expressed in % v/v relative to the same depth, removes the vertical moisture gradient and image-to-image differences in overall wetness, and allows rooted and root-free cells to be compared within the same rhizotron. It was compared among cells without roots and cells with roots of increasing age with a mixed model including treatment, phase and depth, with plant as a random effect. The drying rate of a cell was the change in VWC between consecutive scans divided by the interval in days, analyzed for the deficit period (29-38 DAT) as a function of root age class, root pixel density, cell depth, and starting VWC. The within-cell coupling between brightness and moisture was the change in root brightness between consecutive scans regressed on the change in local VWC, root age, depth, and treatment (a random sample of 120,000 cell pairs). Colonization of root-free cells between consecutive scans was modeled by logistic generalized estimating equations with plant as the cluster, as a function of the cell’s same-depth moisture anomaly, depth, distance (in cells) to the nearest rooted cell and treatment, separately by phase, for cells within six cells of an existing root. Segment brightness was modelled on a random sample of 120,000 segments as a function of segment-support class (reference: support 1), segment age, depth, local VWC, treatment and genotype, with plant as a random effect, with and without segment width (mask pixels per unit length) as a covariate, and the brightness of transport (support ≥ 2) and terminal (support 1) segments was compared within plant-date pairs by a Wilcoxon signed-rank test. To test whether new roots targeted wetter substrate, cells that were root-free at 28 DAT and colonized by 38 DAT were compared with cells that remained root-free in terms of their same-depth moisture anomaly at 28 DAT, averaged per plant.

Broad-sense entry-mean heritability was estimated separately for each imaging day from 10 DAT onward (fewer than 25 plants had roots at 6 DAT) as H^2^ = V_G_/(V_G_ + V_E_/r), where V_G_ and V_E_ are the genotype and residual variance components from a mixed model with treatment as a fixed effect and genotype as a random effect (restricted maximum likelihood), and r is the harmonic mean number of plants per genotype on that day. Confidence intervals were obtained by bootstrapping plants within each genotype (100 replicates). The stability of genotype rankings over time was quantified as the Spearman correlation between treatment-adjusted genotype means on each imaging day and at 42 DAT. Scan-to-scan measurement noise was estimated from consecutive scans (2-3 days apart, starting at 24 DAT) as the standard deviation of the change in each trait after subtracting the mean change for the same genotype on the same day (log ratios for size and count traits); this is an upper bound because it includes true plant-to-plant variation in growth.

## Results

### An accessible hardware and software pipeline for repeated root imaging

The pipeline combines four components (Figure 1). Custom acrylic rhizotrons (Figure 1a) hold 27.3 × 36 cm of substrate behind a transparent front panel, which is protected from light by a removable black ABS plastic cover. A consumer-grade A3 flatbed scanner captures the entire front panel at 300 dpi in about 10 s (Figure 1b). Including handling, one operator scanned the 110 rhizotrons in about 1.5 hours per session, three times per week. BiRefNet, applied zero-shot to the aligned scans, returns a root mask that preserves fine laterals and root tips against the dark, heterogeneous coco coir-peat background (Figure 1c); the only user-set parameters were the binarization threshold and the cleaning rules. A cleaning step removes noise and marks left by earlier experiments, and the cleaned mask is converted into a skeleton graph from which tips, branch points, segment orientations, and segment support are extracted (Figure 1d). Because scans are aligned across dates, the same images also yield a soil moisture map (Figure 1e), a root brightness map (Figure 1f), and depth-resolved root traits (Figure 1g). Across the 1,689 scans, BiRefNet masks required minimal cleaning (median 0.13% of pixels altered), and 86% of the scans of treated plants passed quality control (Figure S2); most exclusions were early scans in which condensation or the absence of roots left nothing to segment.

### Temporal development of root systems differs among rootstocks

Overlaying the aligned masks of all replicates by genotype and imaging day (Figure 2) shows the progressive colonization of the rhizotron and reveals two features of rootstock behavior. Genotypes differed markedly in the timing of root emergence and in root system architecture. Freedom, 101-14, 3309C, and GRN-3 had visible roots in most replicates by 10 DAT, whereas only two of ten 110R and two of nine Ramsey plants had roots at 10 DAT. Most 110R, Ramsey, 5C, and GRN-1 cuttings produced their first visible roots between 12 and 19 DAT (Figure 2, first columns; Figure 3); 1103P rooted at an intermediate time but then grew fastest. Root age maps (Figure 2, last column), in which each pixel is colored by the day the roots first occupied it, show that the earliest roots of all genotypes were long, sparsely branched adventitious roots that reached the bottom of the rhizotron by 28-31 DAT, and that most of the root mass was added later by lateral roots along these axes. Treatment-specific trajectories of total root length are shown in Figure S4.

**Figure 2.**
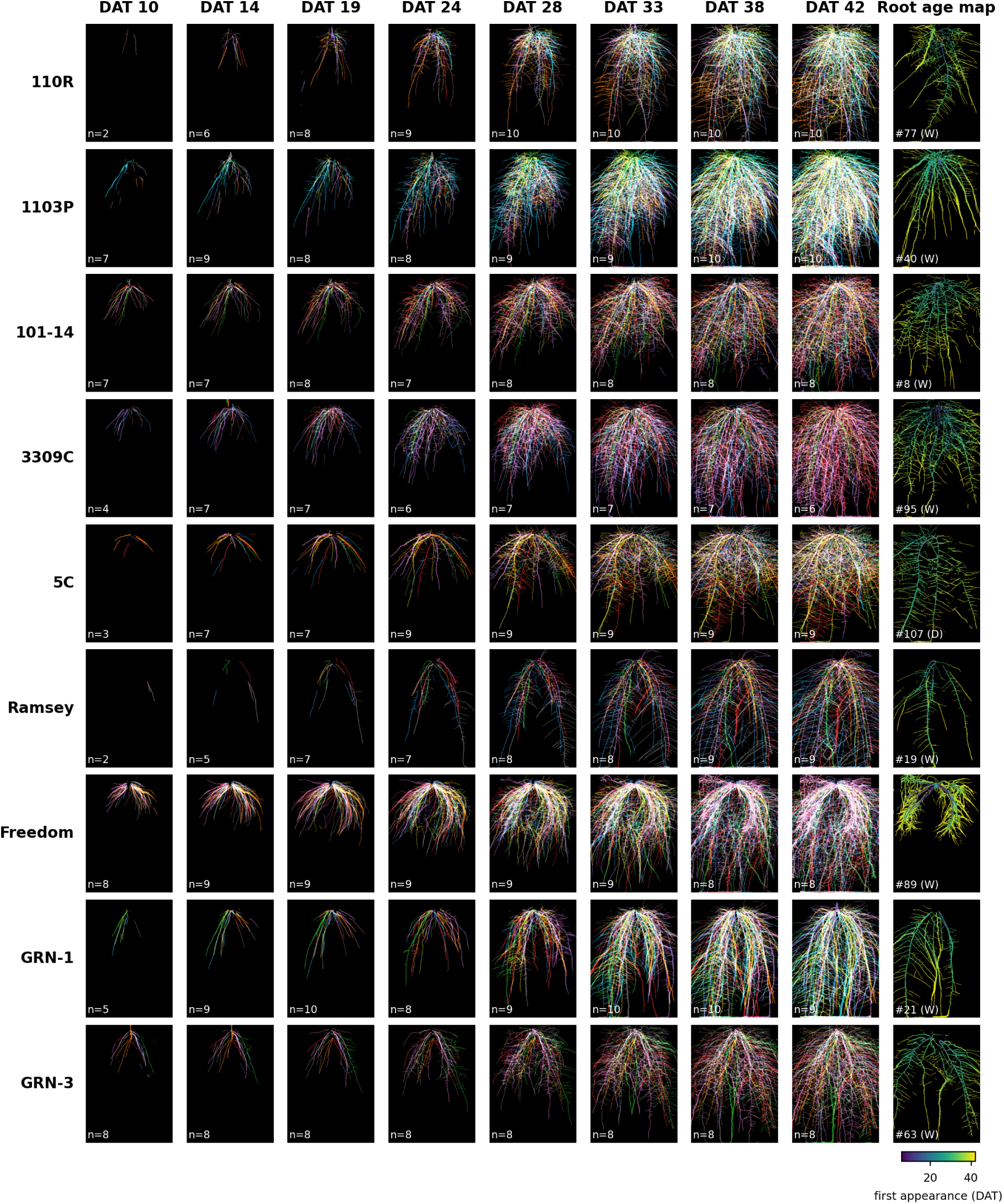
Temporal progression of root systems by genotype. Rows represent genotypes, and columns represent days after transplanting (DAT). In each panel, the aligned, cleaned masks from all analyzed replicates are overlaid, with each replicate shown in a different color; n indicates the number of replicates with a retained image on that day. The last column shows a root age map for one representative plant per genotype (the well-watered plant closest to the genotype median root area at 42 DAT, or any plant when fewer than three well-watered plants were available; W, well-watered; D, water deficit), in which each pixel is colored by the day on which roots first occupied it.

**Figure 3.**
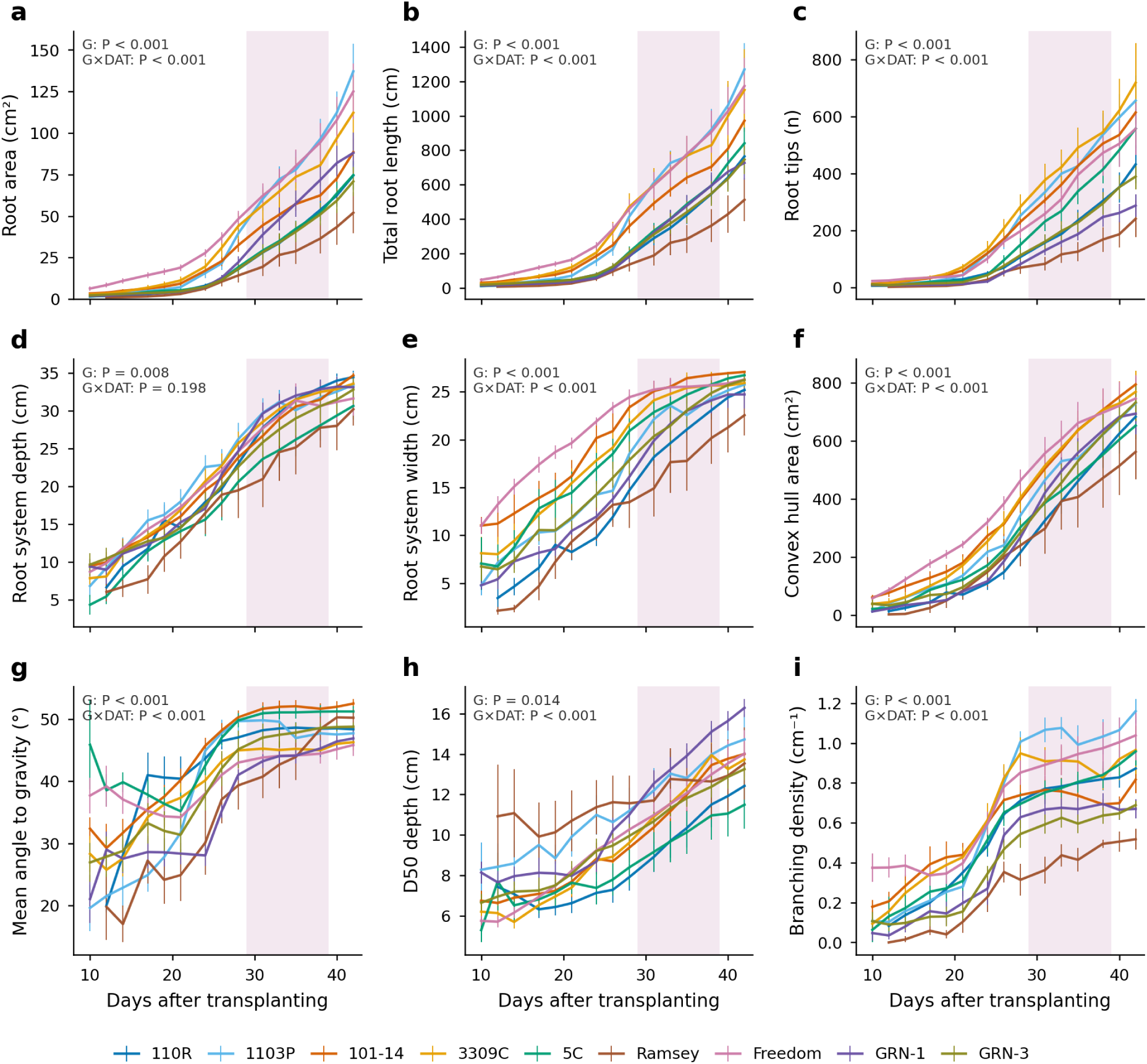
Trajectories of whole-root-system traits. Genotype means ± SE by imaging day from 10 DAT, with treatments pooled (treatment is a covariate in the models), for (a) root area, (b) total root length, (c) number of root tips, (d) root system depth, (e) root system width, (f) convex hull area, (g) length-weighted mean segment angle to gravity (0°, vertical), (h) D50 depth, the depth above which half of the root length lies, and (i) branching density. FDR-adjusted P values for the genotype (G) and genotype × day (G×DAT) effects from the mixed models in Table S2 are shown in each panel. The shaded band marks the water-deficit period (29-38 DAT).

All nine whole-root traits shown in Figure 3 changed significantly with imaging day and, except for root system depth, differed among genotypes in their trajectories (genotype × day interaction, FDR < 0.001; Table S2). Across the full set of 21 traits, the genotype effect was significant for all traits except tortuosity and mean segment support, and the genotype × day interaction was significant for all traits except depth (Table S2). Depth was the exception because the deepest plants of every genotype reached the bottom of the rhizotron from 31 DAT onward, after which the trait stopped being informative. At 42 DAT, 1103P (137 cm^2^ root area, 1,270 cm total length), Freedom (125 cm^2^, 1,173 cm), and 3309C (112 cm^2^, 1,150 cm) had the largest root systems and Ramsey (52 cm^2^, 512 cm) the smallest, with 110R, 5C, GRN-3, GRN-1, and 101-14 intermediate. The number of root tips ranged from 241 (Ramsey) to 718 (3309C), and branching density from 0.52 branch points cm^-1^ (Ramsey) to 1.16 cm^-1^ (1103P). Lateral roots began to emerge between 21 and 26 DAT in all genotypes. Their appearance raised the mean segment angle to gravity from 25-40° to 45-53° (Figure 3g), lowered the mean root diameter (Figure S5i), and lowered the fraction of root length within 30° of vertical (Figure S5g), as expected when near-horizontal laterals are added to initially vertical adventitious axes. At 42 DAT, 101-14, 5C, and Ramsey had the most horizontal root systems (mean angle 50-53°), and Freedom, 3309C, GRN-1, and 1103P the most vertical (46-48°).

### Multivariate structure: developmental time on PC1, root system form on PC2

Principal component analysis of 15 traits placed 65.8% of the variance on PC1 and 10.5% on PC2 (Figure 4a-c). All size and count traits loaded on PC1, together with branching density, transport-root fraction, and mean angle to gravity, and observations were ordered along PC1 by imaging day (Figure 4a). PC1 therefore mainly reflects developmental time and overall size. PC2 was defined by aspect ratio, solidity, D50 depth, and total RLD, and separates deep, narrow, space-filling root systems from wide, shallow ones. Genotype had a significant effect on both axes (P < 0.001), and treatment on PC1, reflecting the larger size of the plants assigned to the deficit. Genotype centroids followed distinct trajectories through the PC1-PC2 plane (Figure 4b): Ramsey and GRN-1 occupied the upper (deep, compact) region, whereas 5C, 101-14, and 3309C occupied the lower (wide, shallow) region throughout the experiment.

**Figure 4.**
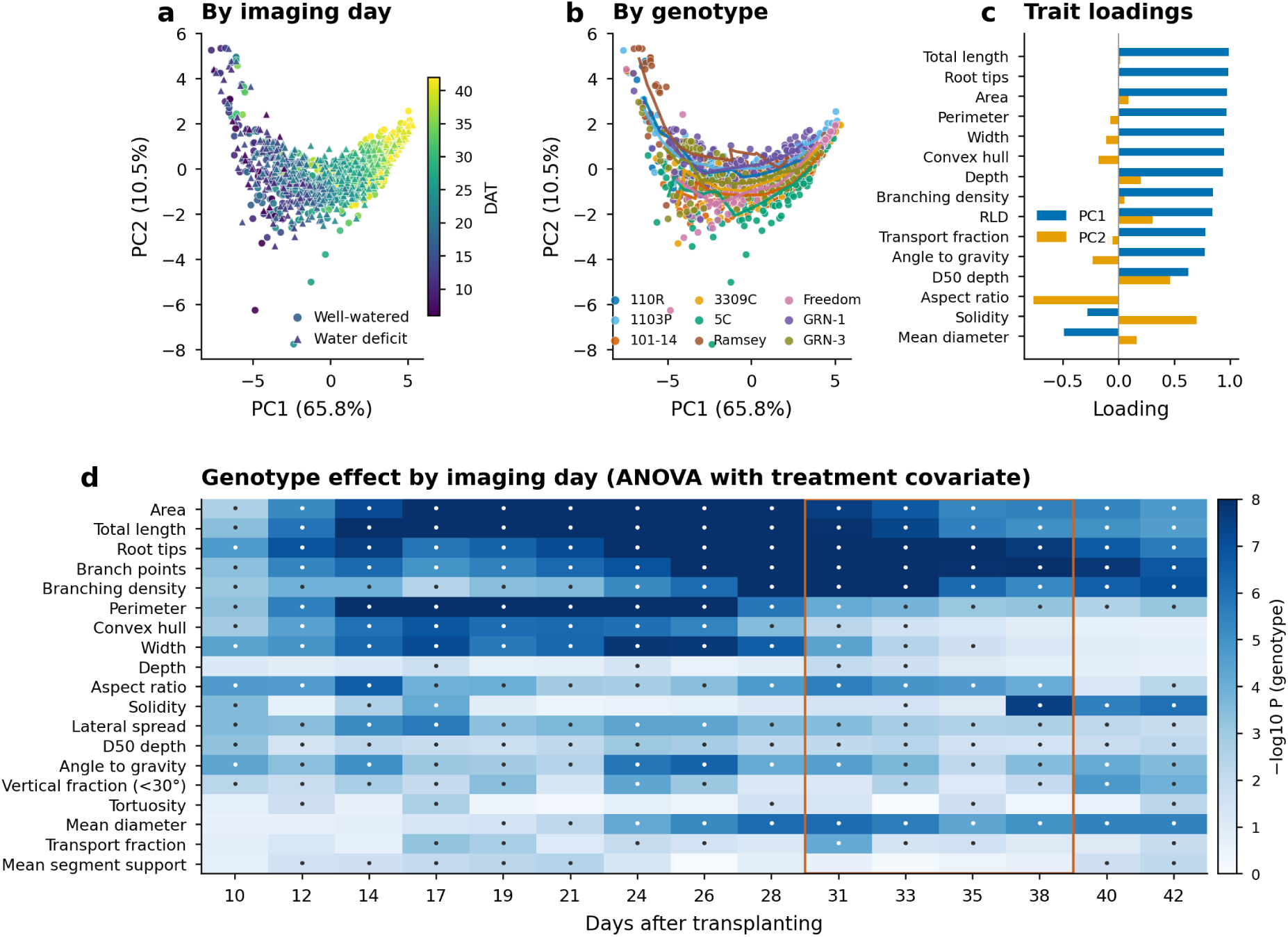
Multivariate structure and the window for genotype discrimination. (a, b) Principal component analysis of 15 scaled traits across 1,108 plant-date observations, colored by imaging day (a; circles, well-watered; triangles, water deficit) and by genotype (b; lines connect genotype centroids in imaging day order). (c) Trait loadings on PC1 and PC2. (d) Strength of the genotype effect for each trait at each imaging day from 10 DAT, expressed as-log_10_ P from an ANOVA with treatment as a covariate. Dots mark trait-day combinations that remain significant after FDR adjustment (FDR < 0.05); the orange rectangle marks the water-deficit period (29-38 DAT).

Testing the genotype effect separately for each imaging day (Figure 4d) identified the window in which the platform discriminates rootstocks best. For root area, the proportion of variance explained by genotype rose from 0.46 at 10 DAT to 0.61-0.63 between 17 and 24 DAT, then declined to 0.35-0.38 from 35 DAT onward as root systems filled the rhizotron. Size and branching traits were significant at every imaging day from 12 DAT onward, whereas root system depth was significant only intermittently and never after 33 DAT. Convex hull area lost significance after 33 DAT and width after 35 DAT. Solidity discriminated genotypes at 10-17 DAT and again from 38 DAT, when genotypes differed in how densely they filled a hull of similar size.

### Spatial deployment of roots across the soil profile

Root length density varied with depth (P < 0.001), and the depth profile differed among genotypes (genotype × bin interaction, P < 0.001; the same held for root pixel density). Before the treatment (24 DAT; Figure 5a), all genotypes concentrated their root length at 4.5-13 cm depth (bins Y2-Y3), but the peak differed by three-to sevenfold between Freedom or 3309C and GRN-1 or Ramsey. By 42 DAT (Figure 5b; treatment comparison in Figure 5c), the profiles had diverged in both shape and magnitude. 1103P, 3309C, Freedom, and 101-14 kept a large share of root length at 4.5-18 cm, 5C had the shallowest profile (D50 = 11.5 cm), and GRN-1, 1103P, and Freedom the deepest (D50 = 14.0-16.3 cm). D50 depth differed among genotypes on every imaging day (Figure 4d), and its trajectories differed as well (Table S2), indicating that vertical deployment is a genotype-specific and dynamic property rather than a by-product of root system size. Treatment also shifted the profile. At the end of the deficit (38 DAT), deficit plants held a larger share of their root length below 18 cm than well-watered plants, and their D50 depth was 4.2 cm deeper before adjusting for plant size (Figure 5c); the size-adjusted analysis of this shift is presented below.

**Figure 5.**
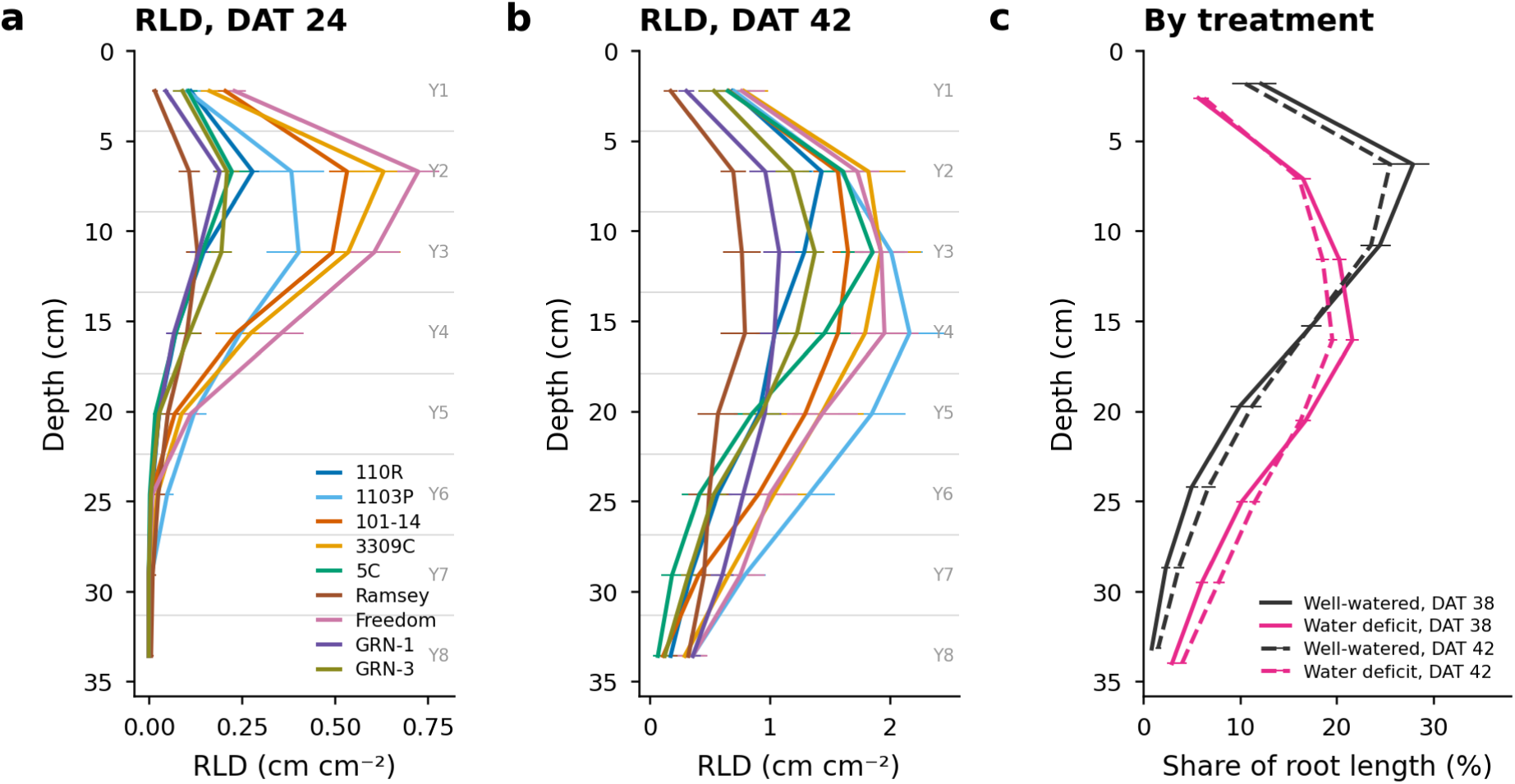
Spatial deployment of roots across the soil profile. Genotype means ± SE of root length density in eight depth bins (Y1-Y8, marked by the horizontal lines) at (a) 24 DAT and (b) 42 DAT. (c) Share of root length by depth in well-watered and water-deficit plants at the end of the deficit (38 DAT, solid) and after recovery (42 DAT, dashed); D50 depth differed between treatments by 4.2 cm at 38 DAT before adjusting for size.

### Root system topology from segment support

Segment support summarizes how many root tips depend on each part of the system (Figure 6a, b). In Ramsey, most tips were served by two or three high-support adventitious axes with short laterals, whereas 1103P at the same age distributed support across many axes and second-order branches (Figure 6a, b). Across genotypes at 38 DAT, 27-36% of root length consisted of segments with support 0, 41-55% of terminal segments (support 1), and 18-25% of segments with support ≥ 2, the transport segments, that is, the shared axes that carry the routes of two or more tips (Figure 6c). Support 0 does not necessarily mean disconnected. About three-quarters of the support-0 length at 38 DAT was connected at both ends but never selected by a route (bridge segments, largely created where roots cross in projection), and the rest was isolated fragments produced by gaps in the mask. Because the support-0 fraction differed among genotypes (24-33% from 24 DAT onward, P < 0.001), we also computed the transport-root fraction over routed length only (support ≥ 1); the two versions were almost perfectly correlated (r = 0.97). The top (Freedom, 3309C, 101-14) and bottom (Ramsey) of the genotype ranking were unchanged, but 1103P, which had the largest support-0 fraction, moved from eighth to fourth (Table S3). The transport-root fraction (the share of root length with support ≥ 2) rose from near zero to about 20% between 14 and 26 DAT, as lateral roots emerged and converted the axes into shared paths, and then stabilized at genotype-specific levels of 18-25% (Figure S5k; Table S2). From 24 DAT onward, Freedom and 3309C had the highest transport-root fractions (24-26%) and Ramsey the lowest (17%; Table S3). Segment orientation also differed among genotypes at 38 DAT. Ramsey and 5C carried the smallest share of root length within 30° of vertical, and 101-14 the largest share of near-horizontal length (Figure 6d). The orientation distributions of well-watered and deficit plants differed only slightly (Figure 6e; change in mean angle 28-38 DAT +1.5° in deficit relative to control plants, P = 0.065).

**Figure 6.**
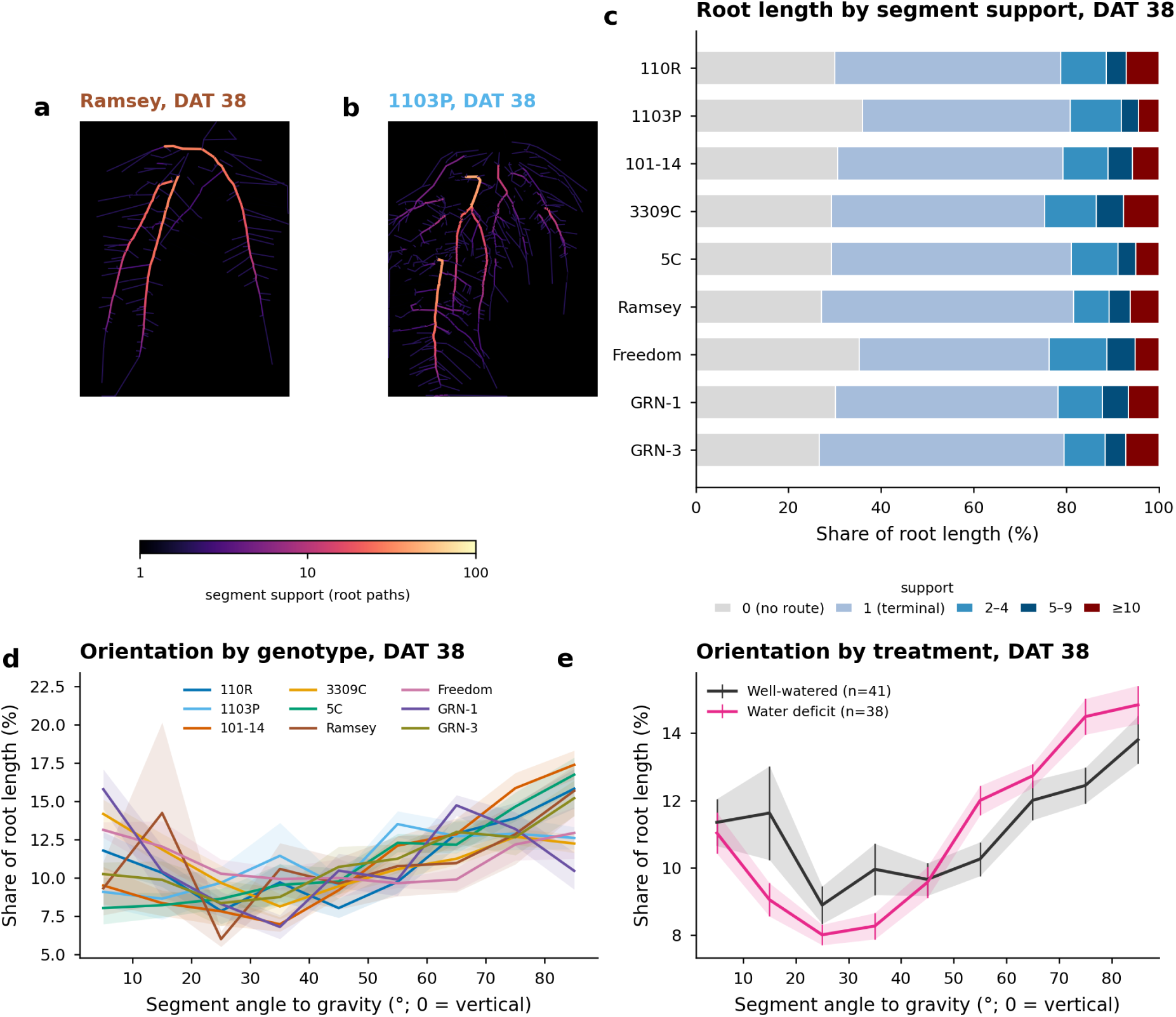
Root system topology from segment support. (a, b) Skeleton segments colored by support (log scale; support 0 drawn as 1) for a Ramsey and a 1103P plant at 38 DAT. (c) Mean share of root length at 38 DAT in five support classes by genotype; support 0 denotes segments not traversed by any route, that is, isolated fragments and connected bridge segments that no route selects (Supplementary Note S1). (d) Length-weighted distribution of segment angle to gravity at 38 DAT by genotype and (e) by treatment (all genotypes); distributions were computed per plant and are shown as means ± SE across plants (bands). The transport-root fraction over time is shown in Figure S5k.

### Image-derived soil moisture tracks the water deficit and its recovery

Root area trajectories of every analyzed plant are shown in Figure 7a; the deficit period (29-38 DAT) is shaded, and the larger starting size of the deficit group is visible. The soil moisture map reproduced the treatment (Figure 7b). Estimated VWC of well-watered and deficit rhizotrons was indistinguishable before 29 DAT (0.44 cm^3^ cm^-3^ in both groups) and diverged during the deficit, reaching 0.31 in deficit rhizotrons at 38 DAT against 0.42 in controls (P < 0.001). After re-watering, VWC of deficit rhizotrons exceeded control values within two days (0.44 versus 0.38 at 40 DAT), with the controls having dried slightly at the end of the experiment. Stomatal conductance (Figure 7c) was not lower in deficit plants than in controls during the first six days of the deficit. It fell below the control value only at 38 DAT, when soil moisture was lowest (0.37 versus 0.46 mol m^-2^ s^-1^; P = 0.17), and rose to 0.51 mol m^-2^ s^-1^ four days after re-watering, when it exceeded the control value of 0.35 (P < 0.001). We attribute the low conductance of the controls to two features of the experiment, which we could not test directly: deficit plants had larger root systems at the start of the treatment, and the controls were irrigated to saturation of a coarse, water-retentive substrate, which may have limited root-zone aeration.

**Figure 7.**
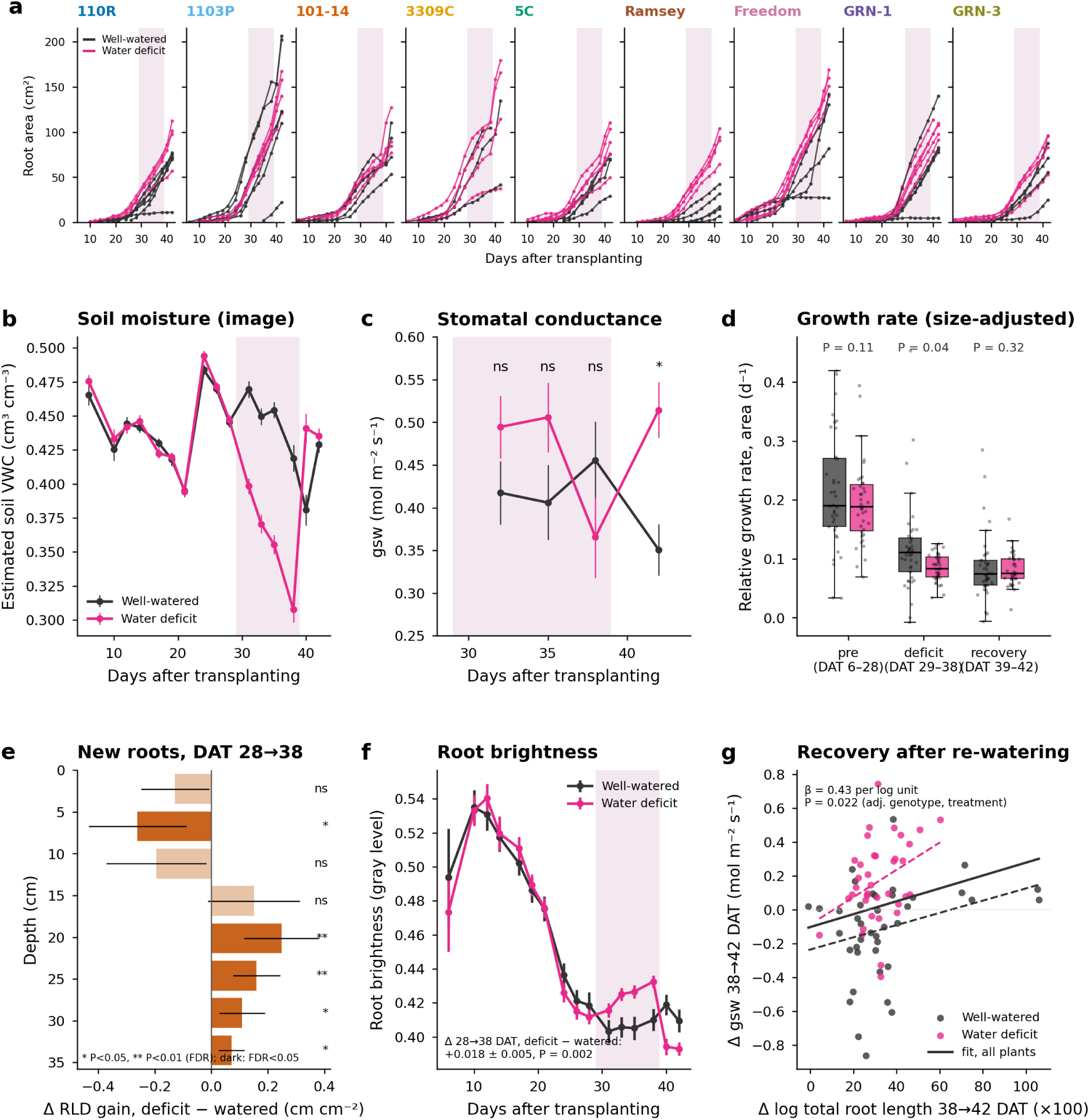
Water deficit, soil moisture, growth, and root brightness. (a) Root area trajectories for each analyzed plant by genotype and treatment; the shaded band marks the deficit period (29-38 DAT). (b) Image-derived volumetric water content (means ± SE) of well-watered and water-deficit rhizotrons over time. (c) Stomatal conductance (means ± SE) at 32, 35, 38, and 42 DAT; asterisks mark dates when treatments differed (P < 0.05; ns, not significant). (d) Relative growth rate of root area per plant and phase; P values are for the treatment effect in size-adjusted mixed models. (e) Adjusted difference (deficit - watered, ± 95% CI) in root length density gained in each depth bin between 28 and 38 DAT; dark bars, FDR < 0.05. (f) Whole-plant root pixel brightness (means ± SE) by treatment over time, with the adjusted 28→38 DAT treatment difference from Table S4. (g) Change in stomatal conductance between 38 and 42 DAT against the change in log total root length over the same interval for individual plants; dashed lines, least-squares fits within each treatment (colored as the points); solid line, fit to all plants.

### Water deficit shifts new root growth to deeper soil without reducing total growth

Because larger root systems were preferentially assigned to the deficit treatment, deficit plants added more root area per day than controls in absolute terms (Figure 7a). After adjusting for size, this difference disappeared. Relative growth rate showed no treatment × phase interaction (P = 0.59), and size-adjusted growth during the deficit was not lower in deficit plants (Figure 7d). No genotype × treatment interaction was detected during the deficit (P = 0.27), but with four to five deficit plants per genotype this test has limited power.

What the deficit changed was where new roots grew (Figure 7e; Table S4). Between 28 and 38 DAT, and after adjusting for baseline values and size, deficit plants added less root length in the upper 13 cm (bins Y1-Y3) and more at 18-36 cm (bins Y5-Y8, all P ≤ 0.010), so that D50 depth increased by 1.3 cm more in deficit plants than in controls (P = 0.002). Deficit plants also produced slightly thinner roots (P = 0.029) and more root tips (P = 0.025), whereas total root length, root area, width, branching density, transport-root fraction, and mean segment angle changed similarly in the two groups (Table S4). The change in root brightness also differed. Roots of deficit plants became brighter (+0.02 gray level, P = 0.002) while those of controls darkened slightly (Figure 7f). Genotype × treatment interactions were not significant for these changes except for aspect ratio and RLD gain in Y2, indicating that the downward shift in new root growth was broadly shared among the nine rootstocks. Restricting the analysis to the 53 plants within the size range shared by both treatments gave the same overall pattern, with less precision (Table S5).

We also asked whether new roots grew preferentially into wetter patches. They did not. At the scale of 0.9 cm cells, the substrate into which new roots grew between 28 and 38 DAT was only marginally wetter than the substrate that remained root-free at the same depth, with no difference between treatments (P = 0.54). In a cell-level model of colonization between consecutive scans, the odds of colonization did not depend on the cell’s moisture anomaly (P = 0.81), but fell by 69% per cell of distance from the nearest root and rose with depth (Table S6). The deeper placement of new roots under deficit is therefore not explained by local targeting of wet patches; it is consistent with the vertical moisture gradient and with growth from existing roots.

### Substrate around roots dries faster the older and denser the roots are

Because every scan is aligned to the same frame, the soil moisture map and the root mask of the same plant can be tracked cell-by-cell over time (Figure 8a). Three simple quantities describe this coupling. The local moisture anomaly is the estimated VWC of a 0.9 × 0.85 cm cell minus the mean VWC of all cells at the same depth in the same image; it is positive where a cell is wetter than its surroundings at that depth and negative where it is drier, and it removes both the vertical moisture gradient and the differences in overall wetness between images, so that cells with and without roots can be compared within the same rhizotron. The root age of a cell is the number of days since roots were first detected in it; “0 (new)” cells were colonized since the previous scan, and cells without roots serve as the reference. The drying rate is the change in a cell’s VWC between two consecutive scans divided by the number of days between them, which measures how quickly water is being removed from that cell. All three were averaged per plant before computing means and standard errors, so the units of replication are plants, not cells.

**Figure 8.**
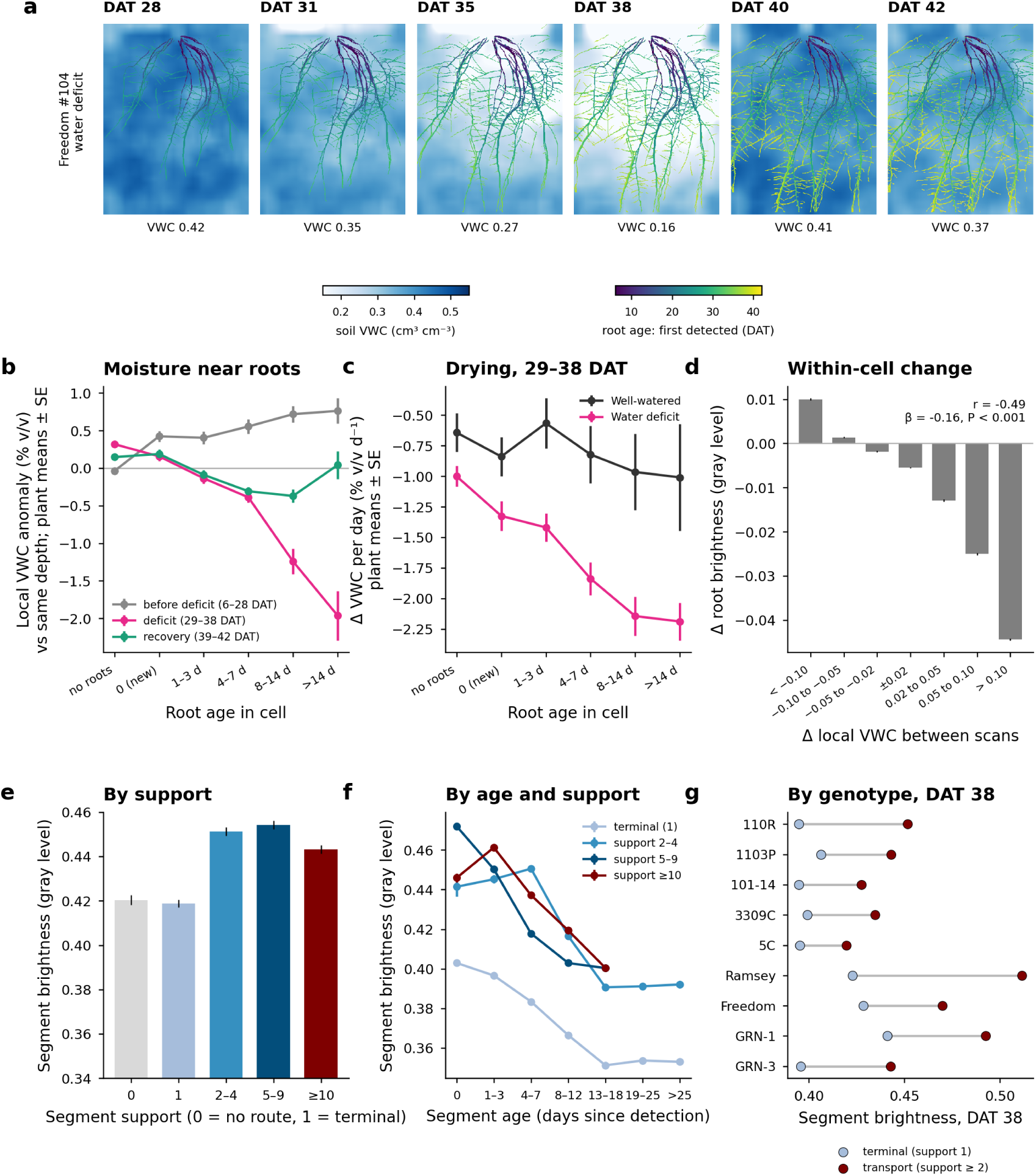
Root-soil moisture coupling and segment brightness. (a) Image-derived soil moisture maps of a water-deficit rhizotron (Freedom #104) from 28 to 42 DAT, with roots overlaid and colored by the day they were first detected; the rhizotron mean VWC is shown in each panel. (b) Local moisture anomaly (cell VWC minus the mean of the same depth row in the same image) for root-free cells and cells with roots of increasing age, before, during, and after the deficit; cell values were first averaged per plant and are shown as plant means ± SE (n = 61-80 plants per point). (c) Drying rate of cells between consecutive scans during the deficit (29-38 DAT) by root age class and treatment (plant means ± SE; n = 34-41 plants). (d) Change in root brightness between consecutive scans as a function of the change in local VWC in the same cell (means ± SE). (e) Mean brightness of skeleton segments by support class (plant-date means ± SE). (f) Segment brightness against segment age by support class. (g) Brightness of terminal (support 1) and transport (support ≥ 2) segments by genotype at 38 DAT.

The aligned moisture maps show the substrate drying from the top and from the root zone outward during the deficit, and re-wetting within two days of irrigation (Figure 8a). Before the deficit, cells containing roots were marginally wetter than root-free cells at the same depth (anomaly +0.4 to +0.7% v/v; Figure 8b), a difference within the range expected from edge and illumination effects (Figure S6c). During the deficit, the pattern reversed and depended on root age. Newly colonized cells were not drier than root-free cells, cells that had contained roots for 1-3 days were only slightly drier, and cells that had contained roots for 4-7, 8-14, and more than 14 days were progressively drier, down to 2.3% v/v below root-free cells for the oldest class (Figure 8b). In other words, at the height of the deficit, the driest patches of substrate were those that had been occupied by roots for a week or more, not the patches that roots had just entered. After re-watering, these differences shrank and largely disappeared (Figure 8b). Within rooted cells, the anomaly also decreased with root density (P < 0.001) independently of age, so the age effect is not just a proxy for the greater root density of cells with older roots.

The drying rate during the deficit showed the same ordering (Figure 8c). In deficit rhizotrons, root-free cells lost 1.0% v/v per day, cells with new roots 1.3%, cells with roots aged 4-7 days 1.8%, and cells with roots older than 8 days 2.1-2.2% per day. In well-watered rhizotrons, which were re-irrigated between scans, drying was slow and did not depend on root age, which confirms that the age gradient in deficit rhizotrons reflects water uptake rather than a property of the imaging. Adjusted for root density, depth, and starting VWC, drying was faster around older roots and slower around brighter roots (both P < 0.001). At the resolution of this analysis (0.9 cm cells, 2-3 day intervals), local water depletion therefore scaled with the amount and age of roots present, not with the presence of young, bright roots.

### Root brightness is related to local soil moisture, root age, and segment support

At the whole-plant level, root brightness declined from 0.53 at 10 DAT to 0.39-0.41 at 42 DAT in both treatments as root systems aged and laterals accumulated. It rose in deficit plants during the deficit and dropped sharply within two days of re-watering (Figure 7f). At the scale of individual grid cells, root brightness decreased with local soil moisture after accounting for root age, cell depth, treatment, genotype, and plant (Figure S6a), and it decreased with root age (Figure S6b). Newly detected roots had a mean gray level of 0.43, which fell to 0.38 within two to two and a half weeks and then stabilized. Part of the moisture-brightness coupling is optical. When the soil moisture estimate excluded a wider margin of substrate around the roots, the slope dropped to about a quarter of its value (Figure S6c), so most of the cell-level coupling comes from substrate within 5 mm of the root, which shares illumination and edge effects with the root pixels. The moisture slope differed among genotypes (P < 0.001), and genotypes also differed in mean brightness (GRN-1 and Freedom 0.44 and Ramsey 0.42, versus 101-14 at 0.39). Within cells followed across consecutive scans, changes in brightness tracked changes in local moisture (r =-0.49, P < 0.001; Figure 8d): root pixels brightened when the surrounding substrate dried and darkened when it was re-wetted.

Segment support added an independent component of brightness variation (Figure 8e-g). Averaged per plant and date, terminal and support-0 segments had a mean gray level of 0.42, whereas transport segments (support ≥ 2) reached 0.44-0.45 (Figure 8e). In a mixed model adjusted for segment age, depth, local VWC, treatment, and genotype, segments with support 2-4, 5-9, and ≥ 10 were 0.054, 0.078, and 0.090 gray levels brighter than terminal segments; the differences shrank but persisted after adding segment width as a covariate and when only thin segments were analyzed (Table S7). Within plant-date pairs, transport segments were on average 0.042 gray levels brighter than terminal segments (P < 0.001), and the difference was present in every genotype at 38 DAT, from 0.025 (5C) to 0.089 (Ramsey) (Figure 8g). Brightness declined with age in terminal segments from the day of detection, whereas transport segments brightened slightly over their first week (as the thin lateral that was first detected became a thicker axis) and then declined, and they remained brighter than terminal segments of the same age (Figure 8f).

### Root system size is associated with stomatal conductance in well-watered plants

After pairing 313 gsw measurements with the nearest root image and adjusting for treatment, genotype, date, and plant, eleven root traits were positively associated with gsw (FDR < 0.05; Figure 9a; Table S8). The strongest associations were for root system width, convex hull area, depth, total root length, root area, and RLD in the upper soil profile. RLD in the lower half of the rhizotron, the number of root tips, D50 depth, solidity, transport-root fraction, mean diameter, root brightness, and image-derived soil moisture were not associated with gsw. These associations were confined to the well-watered group. Within well-watered plants, width and total length had clear effects, whereas within deficit plants no trait was associated with gsw (Figure S7). Adjusting for root area removed all associations except that of width (P = 0.024). The pattern therefore mainly reflects overall root system size, with an additional contribution of lateral extent, among well-watered plants in which the smallest root systems belonged to plants whose growth had stalled (Figure 7a). Within plants, the recovery of gsw after re-watering (38 to 42 DAT) was positively related to the root length added over the same four days (P = 0.022; Figure 7g).

**Figure 9.**
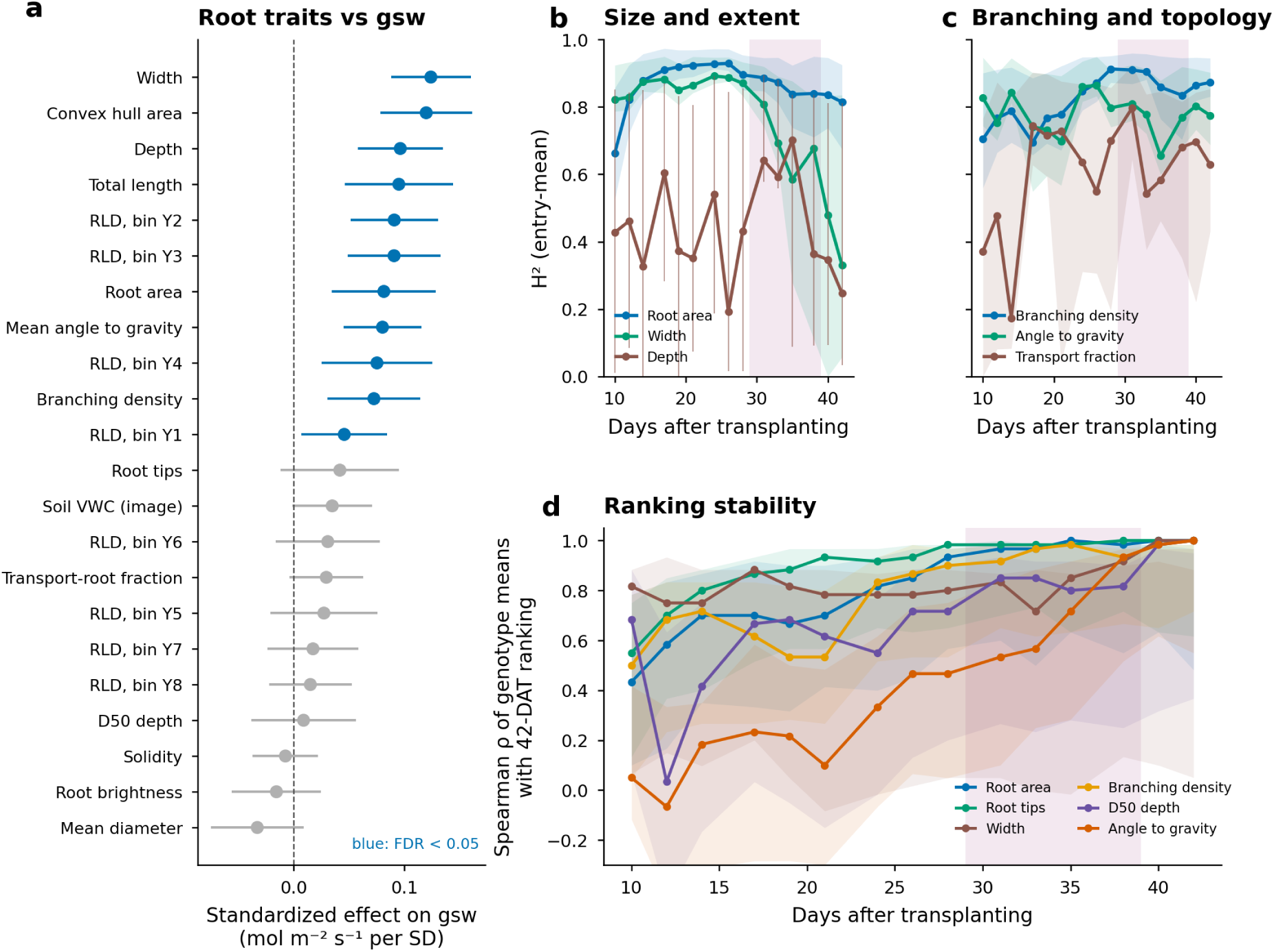
Associations with stomatal conductance and temporal heritability. (a) Standardized effect (± 95% CI) of each root trait on stomatal conductance from mixed models adjusted for treatment, genotype, date, and plant; blue indicates FDR < 0.05 (within-treatment estimates in Figure S7). (b, c) Broad-sense entry-mean heritability of image-derived traits estimated separately for each imaging day, with 95% confidence bands from bootstrapping plants within genotypes (100 replicates); the shaded band marks the water-deficit period. Estimates for all traits are in Figure S8 and Table S9. (d) Spearman correlation between treatment-adjusted genotype means at each imaging day and at 42 DAT, with 95% confidence bands from bootstrapping plants within genotypes (200 replicates).

### The genetic signal in image-derived traits changes with plant age

Entry-mean heritability was estimated separately for each imaging day (Figure 9b, c; Figure S8) and showed that most image-derived traits carried a strong genetic signal, but that the signal changed with plant age in a trait-specific way. Root area and total root length rose from H^2^ = 0.66-0.73 at 10 DAT to 0.93 at 21-26 DAT and declined to 0.81 at 42 DAT; root tip number was at least 0.81 on every day and peaked at 0.92; branching density increased to 0.83-0.91 after 28 DAT; and mean angle to gravity stayed at 0.66-0.86 throughout (Figure 9b, c; Table S9). Traits bounded by the rhizotron behaved differently. Heritability of root system width was 0.82-0.89 until 26 DAT and then fell to 0.33 at 42 DAT, convex hull area fell from 0.87 to 0.25, and root system depth never exceeded 0.70 and fell to 0.25 once all genotypes had reached the bottom. Depth-resolved RLD at 4.5-9 cm (Y2) was among the most heritable traits (0.74-0.94), whereas RLD in Y3 became heritable only after roots reached that layer. The transport-root fraction reached H^2^ = 0.80 at 31 DAT. The genetic share of variance for size traits peaked at about 60% between 19 and 26 DAT (Figure S8), defining the window in which a single scan is most informative for selection in rhizotrons of this size. Genotype rankings converged on their final (42 DAT) order at trait-specific times (Figure 9d). Rankings for root tips and width were already strongly correlated with the final ranking by 14-17 DAT, root area and branching density by 24-28 DAT, and D50 depth by 31 DAT, whereas the ranking for mean angle to gravity did not stabilize until lateral roots dominated the skeleton (after 33 DAT). Scan-to-scan variation between consecutive images, after removing the mean growth of each genotype on each day, was 15% for root area and total length, 22% for tips and 32% for branch points, but only 4-6% of the trait mean for width, depth, D50, mean angle, and diameter, and 3.5% for brightness (Table S10); these values bound the measurement noise of the pipeline and explain why count traits are noisier than length and extent traits.

## Discussion

### A zero-shot segmentation model removes the annotation bottleneck of rhizotron phenotyping

The main practical result of this study is that a general-purpose segmentation network, applied without any training on root images, produced masks of grapevine root systems in a dark, heterogeneous organic substrate that required almost no correction and supported quantitative analysis of 21 traits across 1,108 images. Earlier root image analysis tools either require manual tracing (SmartRoot, RootNav), task-specific training with annotated images (RootPainter), or degrade on complex, overlapping root systems [21]. To our knowledge, this is the first application of a dichotomous image segmentation model to root images; recent systematic comparisons of root segmentation methods [42] evaluated U-Net variants, DeepLab, RootPainter, and fine-tuned SAM-family models, all trained on annotated root images, whereas BiRefNet was used here without any root training. BiRefNet was designed for high-resolution objects with thin structures and complex boundaries, which is precisely the geometry of a root system on a rhizotron panel, and it preserved fine laterals that a stricter threshold removed. The cost of the approach is the GPU time needed for inference on 14-megapixel scans, which can be rented on demand, and the fact that the model segments any bright, thin object; the marks left by roots of earlier experiments on four rhizotrons were the only systematic false positives and were removed by a simple spatial rule; scanning each substrate-filled rhizotron before transplanting would also provide a root-free background image that could be subtracted to suppress such marks. We anticipate that fine-tuning BiRefNet on a small set of root masks would further improve consistency across dates, but the zero-shot results were sufficient for every analysis reported here.

The hardware side of the pipeline is deliberately simple. Acrylic rhizotrons can be built in a laboratory at low cost, and an A3 flatbed scanner costing a few hundred dollars produces 300 dpi images in 10 s, enabling one person to scan 110 rhizotrons three times per week. The same scanner resolves root hairs at 1,600 dpi, a trait that has not been quantified in grapevine root systems, though at the cost of longer scan times. Compared with camera-based rhizotron systems [10] and the luminescence-based GLO-Roots platform [32], the flatbed scanner provides uniform illumination and geometry without a dark room or a custom imaging station, which simplifies alignment across dates and makes the soil moisture mapping possible.

### Root traits resolve rootstock identity and its temporal dynamics

The nine rootstocks differed in nearly every trait examined and, more importantly, in how those traits changed over time. Two groups emerged. Freedom, 3309C, 101-14, and 1103P developed large, highly branched systems, whereas Ramsey, GRN-1, GRN-3, and 110R remained smaller and less branched throughout the six weeks. Ramsey and 110R were also the slowest to produce visible roots, whereas GRN-3 rooted early but grew slowly. The ranking of 1103P and Freedom above 110R and Ramsey in root system size at this stage aligns with their reputation as vigorous rootstocks and with the difficulty of rooting Ramsey and *V. champinii*-derived material from dormant cuttings. Beyond size, the platform resolved differences in form. Aspect ratio, solidity, and D50 depth defined the second principal component and separated deep, compact systems (Ramsey, GRN-1) from wide, shallow ones (5C, 101-14, 3309C) throughout the experiment. This structure closely resembles that reported for an automated rhizotron platform in Arabidopsis, in which time dominated PC1 and root system shape distinguished accessions along PC2 [9], suggesting that developmental time and root system form are general, separable axes of variation in root phenotyping data.

Genotype-specific vertical deployment was evident both as a genotype × depth interaction in RLD and as genotype differences in D50 depth. Fichtl et al. [43] digitized excavated root systems of the rootstocks 101-14 Mgt, SO4, and 110R in three dimensions and found genotype-specific architectures that changed with vine age. A minirhizotron study in a mature vineyard found that Ramsey concentrated new roots below 20 cm, whereas 140Ru spread them evenly [44]. In our study, 5C and 101-14 were the shallowest genotypes and GRN-1, 1103P, and Freedom the deepest, with Ramsey combining a small, sparsely branched system with an intermediate D50. Genotype-specific vertical distribution therefore appears within four weeks of rooting in a rhizotron and again in mature field vines, suggesting that it is expressed early and is not specific to one phenotyping method, although in the field rooting depth is also strongly shaped by soil and management [17]. An advantage of the rhizotron is that it resolves this deployment non-destructively and continuously from the onset of root development.

### Segment support as an annotation-free description of topology

Root orders cannot be reliably assigned from a two-dimensional mask without manual annotation. There is a rich and valuable body of work on root nomenclature and classification, from architectural taxonomies of root types [45] to community standards for classifying, sampling, and measuring roots [46]. These frameworks are the natural way to describe a root system when individual roots can be identified, but applying them to images requires each root to be recognized and labeled, which in practice means manual annotation. In grapevine, for example, complete root-system topologies have been obtained by excavating root systems and digitizing each adventitious root and its lateral orders point by point [43]; this delivers exact root types and orders, but it is laborious and cannot be repeated over time on the same plant. The segment-support metric sidesteps this limitation by asking, for each part of the skeleton, how many root tips depend on it. Segments with high support are the transport axes of the system; segments with support 1 are its absorptive periphery. The metric has three useful properties. It requires no training data. It describes position and function along a root rather than order, so that the base, middle, and tip of a single adventitious root receive different values. And it is a per-segment quantity from which distributions and fractions can be computed for any subset of the image. Its limitation is that it inherits the imperfections of the mask and of the routing rules. A piece of root separated from the rest of the system by a mask gap, for example a lateral whose junction with the root it branches from is hidden by substrate, becomes an isolated fragment, and where roots cross in projection, skeletonization creates short connected bridge segments that no route selects; both receive support 0, and together they accounted for 27-36% of root length, a fraction that differed among genotypes. Support-0 segments keep their length and orientation, and the extremes of the genotype ranking of the transport-root fraction were unchanged when it was computed over routed length only, but tip counts are inflated by the free ends that gaps create and by the crossing artifacts of skeletonization. Temporal information could be used to repair these gaps, since a root that is disconnected on one date is usually connected on an earlier or later one, and we see this as the most valuable next improvement to the pipeline.

The transport-root fraction rose sharply between 14 and 26 DAT and then plateaued at genotype-specific values of 18-25%, reflecting the transition from a system of adventitious axes to a branched system in which the axes become shared paths. Freedom and 3309C invested the largest share of root length in transport segments, whereas 1103P, Ramsey, and 5C invested the smallest, for different reasons: 1103P because of its very large periphery of terminal laterals, and Ramsey because it had few laterals at all. Whether the transport-root fraction relates to hydraulic conductance or to the “coarse root” fraction obtained from excavated systems [47] remains to be tested, but it provides a quantitative, heritable descriptor of branching organization that is not available from conventional traits.

### Water deficit relocates root growth without a detectable reduction in size-adjusted growth

The water-deficit experiment was affected by two design decisions that we discuss openly because they matter for interpretation. Deficit was imposed only on plants with an established root system, so deficit plants were larger than controls at the start of the treatment; and the well-watered controls were irrigated frequently to saturation of a coarse coir-peat substrate, which likely reduced oxygen availability in the root zone. Both factors are visible in the data. Absolute growth rates were higher in deficit plants before the treatment began, stomatal conductance was higher in deficit plants at the first measurement, and several well-watered plants stopped growing altogether between 24 and 42 DAT (Figure 7a). We therefore based all treatment inferences on within-plant changes adjusted for baseline size, and we caution that the “well-watered” group represents a wet rather than an optimal control.

With those adjustments, we found no evidence that the ten-day deficit reduced the root system’s relative growth rate, but the deficit changed where growth occurred, and this shift survived restriction to plants of overlapping size. Deficit plants added less root length in the upper 13 cm and more in the lower 18 cm of the rhizotron, deepening D50 by 1.3 cm more than controls over ten days, and they produced thinner roots and more tips. This is the response expected from the soil moisture maps, which showed the substrate drying from the surface downward and from the root zone outward (Figure 8a), and it resembles the deeper root deployment under drying reported for rootstocks in the field [13,14], although at the centimeter scale growth was not directed toward wetter patches. No genotype × treatment interaction was detectable for most traits, but with four to five deficit plants per genotype only large interactions could have been detected; genotype differences in drought response may require more replication, a longer or more severe deficit, or traits that this two-dimensional system cannot capture, such as root hydraulic properties. The rapid recovery of stomatal conductance after re-watering, and the burst of root growth in deficit plants after 39 DAT, are consistent with the rewetting response attributed to drought-tolerant rootstocks, in which rapid re-establishment of root growth and of water uptake near the root tips after re-watering distinguishes resistant from sensitive genotypes [19].

The image-derived soil moisture estimate performed well as a treatment covariate and as a spatial map, but it is a calibration based on the gray level of a single substrate under a single scanner and should be re-derived for other substrates and imaging set-ups. Its main value in a breeding context is that it costs nothing beyond the scan and provides a within-rhizotron moisture field against which root placement can be evaluated, as done here at the cell level.

### Root brightness, root age, and water uptake

Root pixel brightness carried information at several scales. It decreased with root age at the cell and segment levels; it decreased with local soil moisture and tracked moisture changes within cells; it increased in whole plants under deficit; and it was higher on high-support transport axes than on terminal laterals. A decline in brightness with age is expected from the suberization of grapevine fine roots [20,48], and the stabilization after two to three weeks suggests that the transition from white, absorptive roots to brown, suberized roots can be timed from rhizotron images. The white fraction of the root system is itself meaningful; under water stress, the drought-resistant rootstock 110R kept a larger share of its fine-root length white and functional than the drought-sensitive 101-14 [18], and the brightening of deficit root systems observed here, together with their extra root tips, may reflect the same maintenance of young absorptive roots. The higher brightness of transport axes, which persisted after adjusting for age, width, and local moisture and was observed in all nine genotypes, is more surprising, because these are the oldest parts of the system. Adventitious roots of grapevine cuttings are thick and white with a large cortex, whereas first-order laterals are thin and turn brown quickly; the difference may therefore reflect root type rather than age, and it implies that brightness should not be read as a simple proxy for age across root classes. Genotype differences in brightness were consistent across root classes and are candidates for anatomical validation, for which the fine-root anatomy methods already applied to grapevine rootstocks [18,19] provide a direct template.

We hypothesized that young, bright roots would be the primary sites of water uptake and that the substrate around them would dry fastest. The cell-level analysis did not support this at the available resolution. During the deficit, the substrate around newly colonized cells was no drier than the root-free substrate at the same depth, and both the moisture anomaly and the drying rate increased with root age and root density, so the driest cells were those that had contained roots for more than a week. Several factors can explain this. Uptake per unit length may well be highest in young roots, but cells with older roots contain more root length, including laterals that emerged after the cell was first colonized, so uptake per cell scales with root density; depletion is cumulative and persists where it began. The 2-3 day interval between scans also cannot resolve the diurnal dynamics of uptake by root tips. Resolving uptake per unit root length and age will require finer cells, daily scans, and, ideally, independent measurement of local water content. The effect of moisture on brightness is partly optical, because a wet substrate is darker and reduces edge contrast. Excluding a wider margin of substrate around the roots reduced the cell-level slope by about three-quarters, which shows that most of the coupling arises from substrate immediately adjacent to the root; the residual slope at 5-10 mm, the reversal on re-wetting, and the darkening of whole root systems within two days of re-watering are consistent with a change in root surface reflectance, but an optical origin cannot be excluded and anatomical validation is needed.

### Associations with shoot physiology

Root system width, convex hull area, total length, and RLD in the upper 18 cm were positively associated with stomatal conductance after adjusting for treatment, genotype, date, and plant. However, these associations were confined to the well-watered group and, with the exception of width, disappeared when root area was included as a covariate. Among the well-watered plants, those with larger root systems sustained higher stomatal conductance, whereas the plants with the smallest root systems were those whose growth had stalled in saturated substrate; the deficit plants, all of which had established root systems, showed no relationship between root traits and gsw. Alonso-Forn et al. [47] found that rootstocks with higher root length density and a larger proportion of fine roots maintained higher stem water potential under severe deficit, and stem water potential and stomatal conductance are closely related [49]; our data are consistent with such a relationship but cannot separate it from the effect of plant condition in the wet control. The within-plant analysis is more informative. New root growth is the main way grapevines regain root-system hydraulic conductivity [20], and recovery after drought has been linked to renewed elongation and water uptake at the root tips [19]. After re-watering, the recovery of stomatal conductance was correlated with the root length added over the same four days, which is consistent with, but does not demonstrate, such a role of new roots in restoring gas exchange. These associations are correlational, were estimated in a small number of plants, and should be read as a demonstration that image-derived root traits carry physiological information rather than as evidence of specific causal links.

### Heritability changes with plant age and rhizotron size

Estimating heritability on each imaging day, rather than once, revealed that the genetic signal in image-derived root traits is strong but time-dependent. Size and count traits reached H^2^ above 0.9 between 21 and 31 DAT, comparable to or higher than the values reported for root traits in grapevine mapping and diversity populations (0.52-0.70 [50]; 0.36-0.82 [51]); a single destructive measurement of adventitious root traits in a 110R × 101-14 cross gave lower values still (0.08-0.34), with genotype values that did not carry over between propagation methods [52]. However, the heritability of traits bounded by the rhizotron (width, convex hull area, depth) collapsed once roots reached the walls. These estimates are based on nine genotypes and should be read as the genetic signal in this panel rather than as population parameters. Root system depth is the clearest case: its heritability never exceeded 0.70 and fell to 0.25 by 42 DAT because all genotypes reached the bottom. These patterns have a direct practical consequence. For grapevine cuttings in rhizotrons of this size, a single scan between about 20 and 30 DAT captures most of the genetic variance in size, branching, and upper-profile RLD, whereas depth-related traits require either taller rhizotrons or earlier measurement. More generally, they show that a single time-point measurement can under-or overestimate the genetic signal several-fold depending on when it is taken, an argument for time-resolved phenotyping even when only one time point will be used for selection. Repeating the experiment in rhizotrons of different sizes, and tracking when each trait begins to saturate, would test how far these relationships carry over to larger containers and would justify small-container screening as long as measurement stops before the walls are reached. Because root system architecture in grapevine is under complex, multigenic control [52], selection on these traits will likely rely on genomic prediction trained on large phenotyped populations, and image-based pipelines of this kind provide the throughput needed to evaluate the root-system ideotypes that have been proposed for drought-tolerant rootstocks [53].

### Limitations and future work

The pipeline images the roots that grow against the front panel of a 2 cm thick rhizotron and therefore describes a two-dimensional projection of a subset of the root system; roots that grow away from the panel are missed, and the fraction missed may differ among genotypes, so heritability estimates include any genotype-specific bias in what the panel shows. Root tips and branch points are inflated by mask gaps and by crossing artifacts of skeletonization, and we recommend using them as relative rather than absolute counts; the support-0 fraction itself differed among genotypes, so topological traits should be reported over routed length as well as total length, as done here for the transport-root fraction. The water-deficit component of the experiment was limited by assigning larger plants to the deficit and by using a wet control; a follow-up experiment should randomize treatments within size classes at a fixed root size, monitor substrate water content in controls, and extend the deficit. The soil moisture calibration and the interpretation of root brightness require independent validation. Finally, the present analysis used a zero-shot model; fine-tuning on a modest set of curated masks and using temporal consistency across aligned scans to repair gaps and reject false positives are the two most promising improvements to the software. In our view, these limitations do not affect the central conclusion that a scanner, hand-built rhizotrons, and a general-purpose segmentation model can deliver heritable, time-resolved, and depth-resolved root phenotypes for grapevine rootstock breeding.

## Data Availability

Root masks, cleaned masks, extracted traits (per image, per depth bin and per skeleton segment), soil moisture grids, and stomatal conductance data are available at https://github.com/diazgarcialab/grapevine_r oot_architecture_Munoz2026.

## Author Contributions

J.R.M. designed and conducted the greenhouse experiment, built the rhizotrons, collected scans and physiological data, developed the soil moisture calibration, performed data analysis, and drafted the manuscript. E.T.-L. developed the image alignment and BiRefNet segmentation workflow and the segment-support analysis. S.S. and Y.L. contributed to data collection and analysis. A.M. contributed with data analysis and interpretation. L.D.-G. conceived the study, supervised the work, performed the final data analysis, and wrote the manuscript with input from all authors.

## Acknowledgments

The authors thank Veronica Nunez, Ana Gaspar, and Guillermo Garcia Zamora for their support with greenhouse experiments and vineyard maintenance. Artificial intelligence tools were used to write code for the data analysis and figure construction and to improve grammar; all analyses, figures, and text were reviewed and validated by the (human) authors. The authors declare that they have no competing interests.

## Supplementary Material

**Figure S1.**
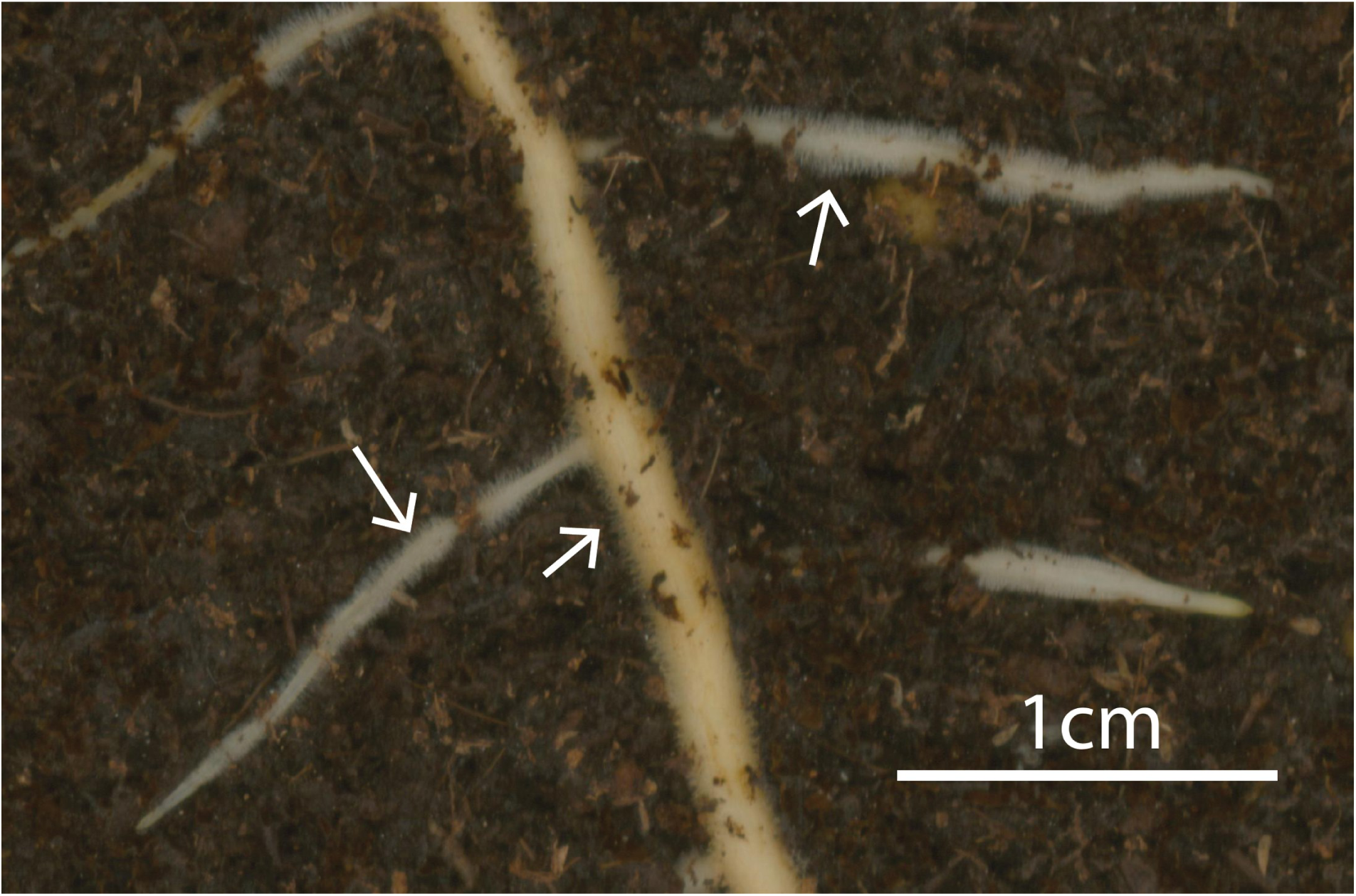
Root hairs resolved by the flatbed scanner at 1,600 dpi. Detail of a rhizotron scan acquired at 1,600 dpi; arrows mark root hairs along young lateral roots and an adventitious axis. Scale bar, 1 cm.

**Figure S2.**
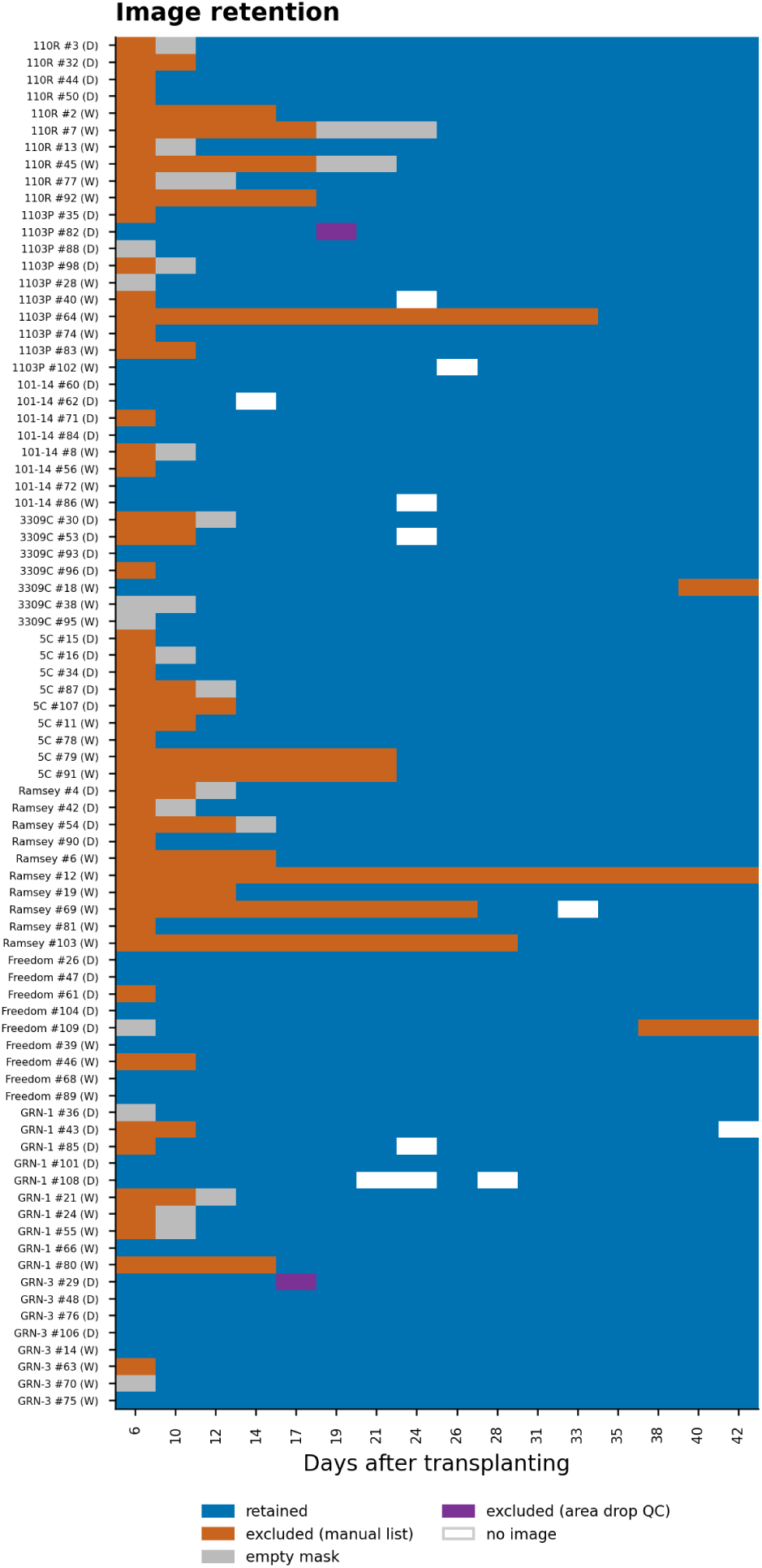
Image quality control. Retention status of every scan of the 85 plants assigned to a treatment (rows) at each imaging day (columns).

**Figure S3.**
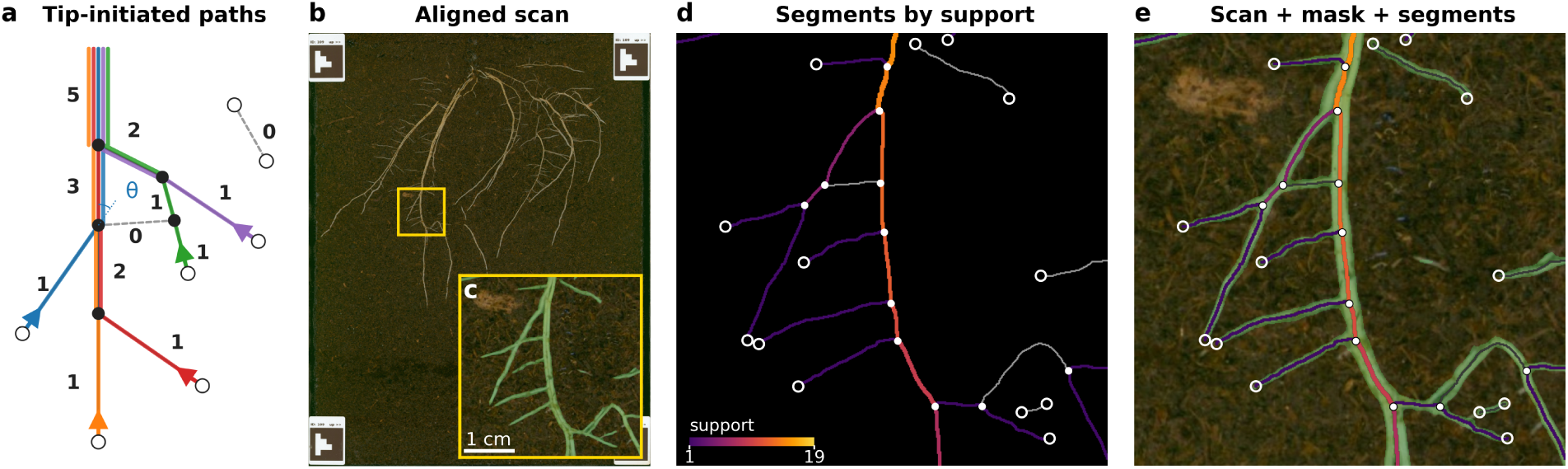
Segment support: construction and a real example. (a) Schematic of the construction. Grey lines are skeleton segments, open circles tips (skeleton nodes of degree one) and filled circles vertices (branch points); the axis continues beyond the frame toward the cutting. Each colored line is one path traced from a tip toward the cutting; arrowheads mark the direction of tracing. At every branch point a path continues along the segment with the smallest turning angle θ relative to its direction of travel. Numbers give the support s of each segment. Dashed segments have support 0: the bridge joining two branch points, created where two roots cross in projection, is connected at both ends but is never the smallest turn for any path, and the isolated segment at the top right, with two tips, starts no path. (b-e) The construction applied to a Freedom rhizotron (ArUco 109) at 24 DAT, before the water-deficit period. (b) Aligned scan; the square marks a 3.8 × 3.8 cm window. (c) The window with the zero-shot BiRefNet mask (threshold 0.3) overlaid in green. (d) Skeleton segments of the window colored by support on a logarithmic scale (support 0 in grey), with line width increasing with support. (e) Scan, mask and segments overlaid.

**Figure S4.**
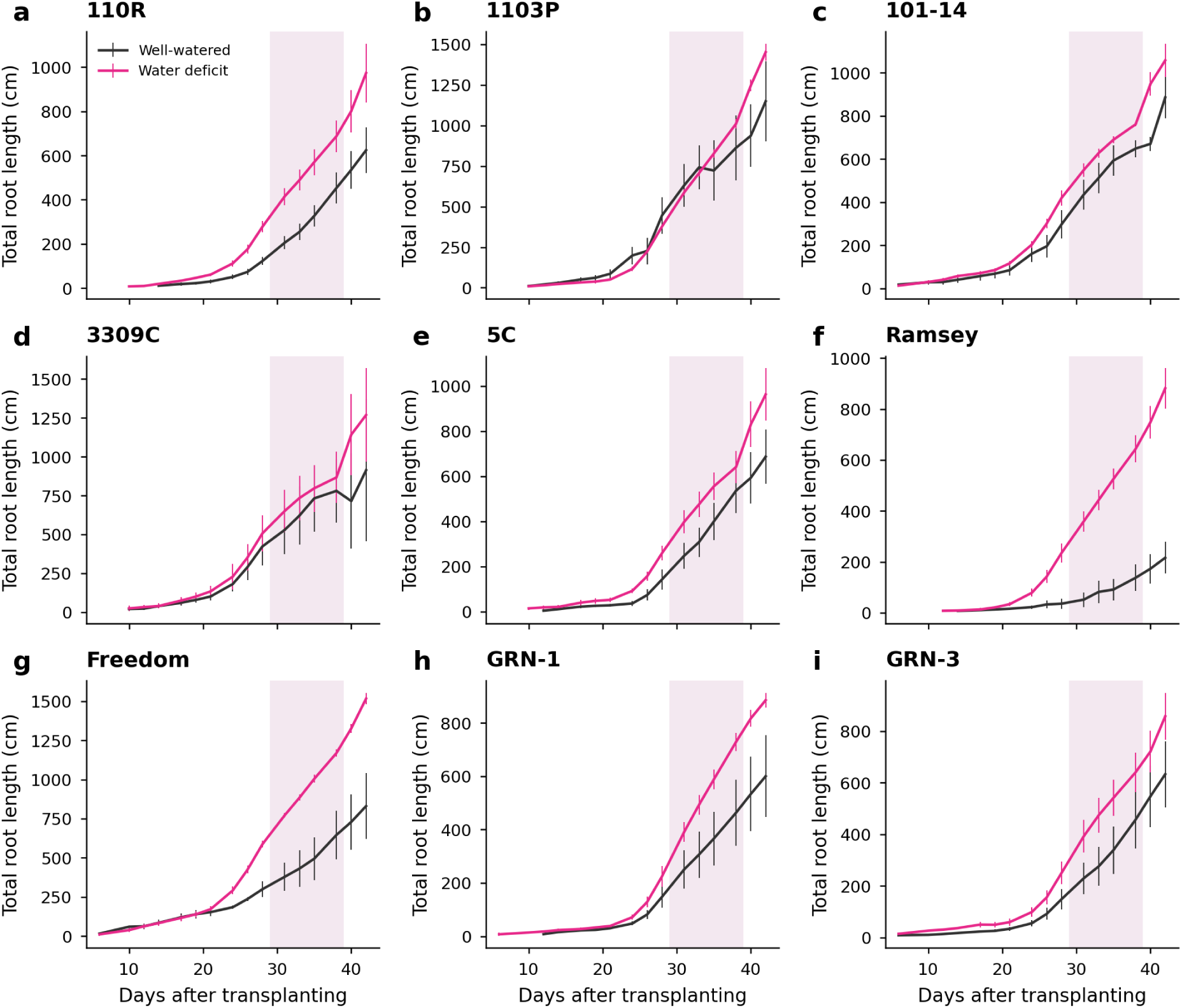
Total root length by genotype and treatment. Means ± SE of well-watered and water-deficit plants for each genotype; the shaded band marks the deficit period. Note that deficit plants were larger at the start of the treatment by design.

**Figure S5.**
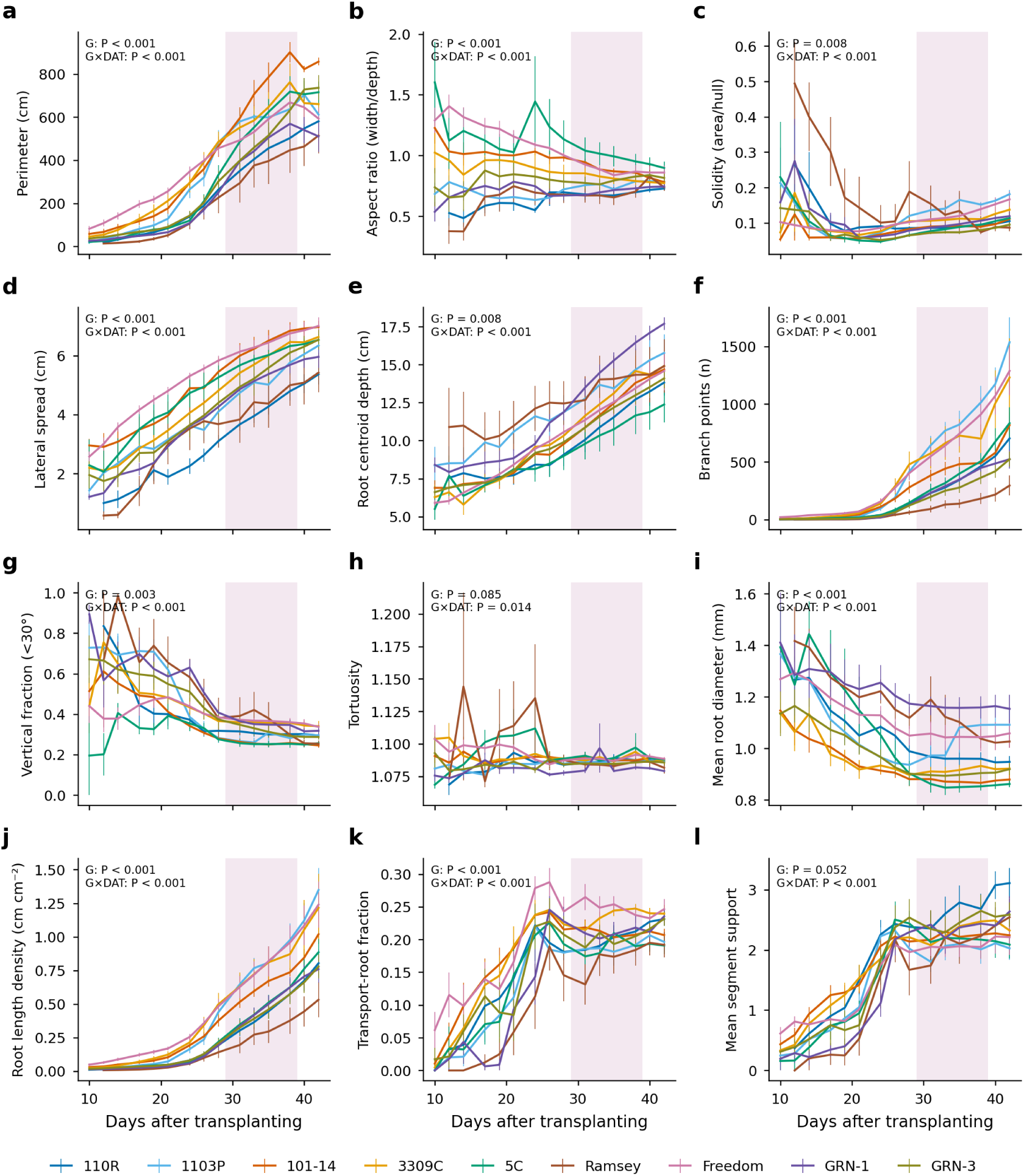
Trajectories of additional traits. Genotype means ± SE by imaging day for (a) perimeter, (b) aspect ratio, (c) solidity, (d) lateral spread, (e) root centroid depth, (f) number of branch points, (g) vertical fraction, (h) tortuosity, (i) mean root diameter, (j) total root length density, (k) transport-root fraction and (l) mean segment support, with FDR-adjusted P values from Table S2.

**Figure S6.**
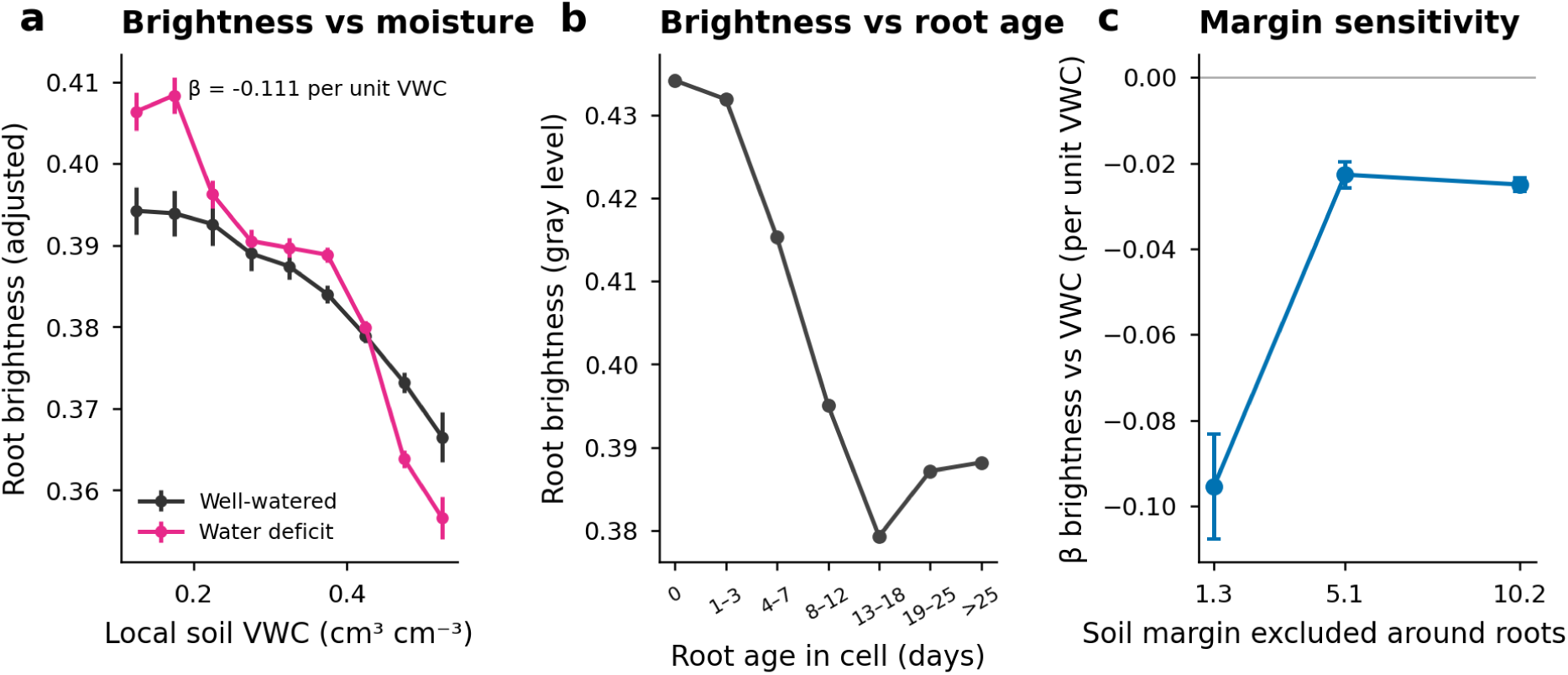
Cell-level root brightness. (a) Root pixel brightness against local soil moisture at the grid-cell level, by treatment, after adjustment for root age, depth, genotype and plant (partial residuals). (b) Root pixel brightness against the number of days since roots were first detected in the cell (means ± SE). (c) Slope of brightness on local VWC when the soil moisture estimate excludes a margin of 1.3, 5 or 10 mm of substrate around the roots (mixed models with plant and plant × date random intercepts; ± 95% CI).

**Figure S7.**
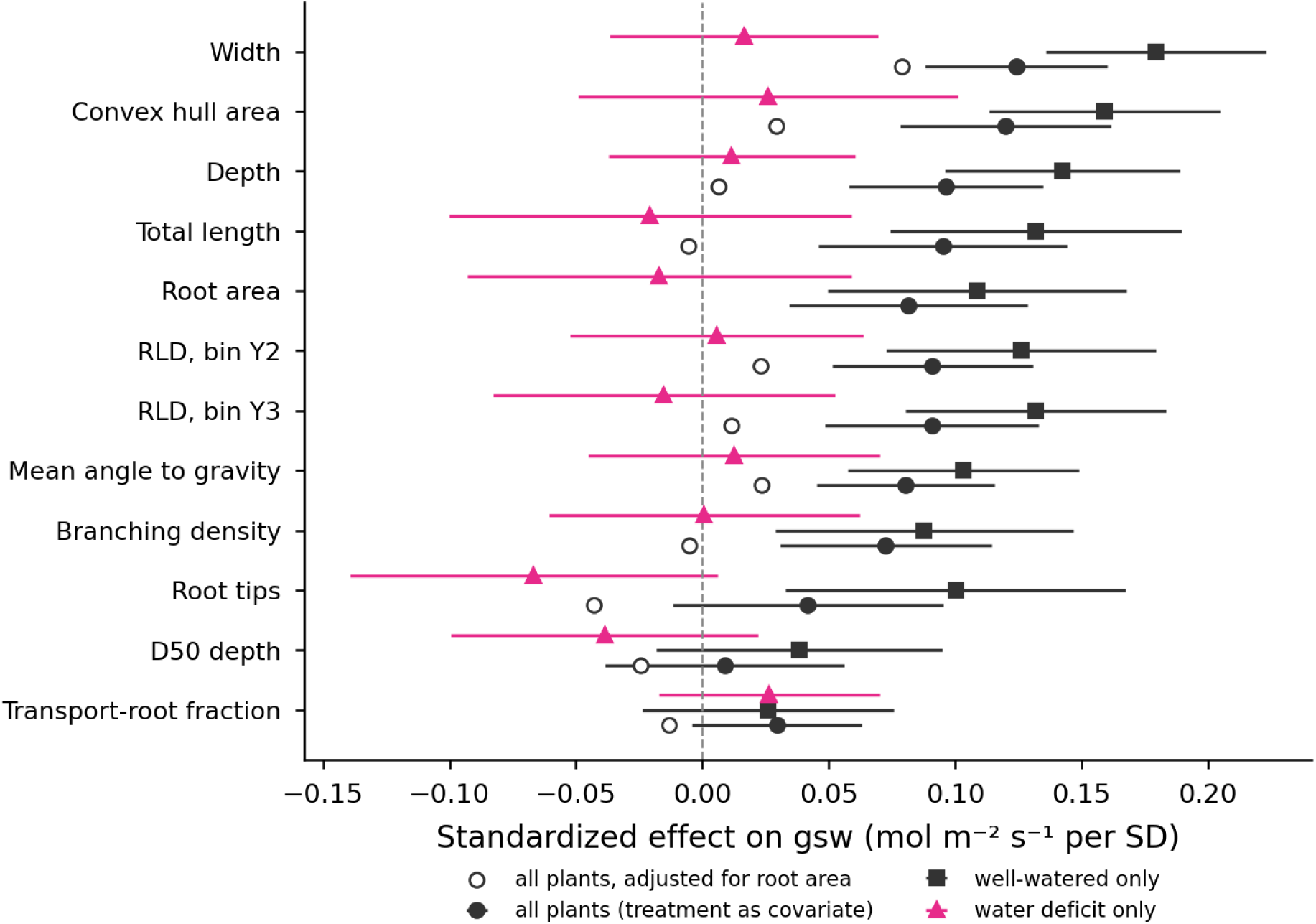
Root trait-stomatal conductance associations within treatment groups. Standardized effects (± 95% CI) of twelve root traits on gsw estimated in all plants with treatment as a covariate (filled circles), in well-watered plants only (squares) and in water-deficit plants only (triangles); open circles are point estimates from the all-plant model with log root area as an additional covariate.

**Figure S8.**
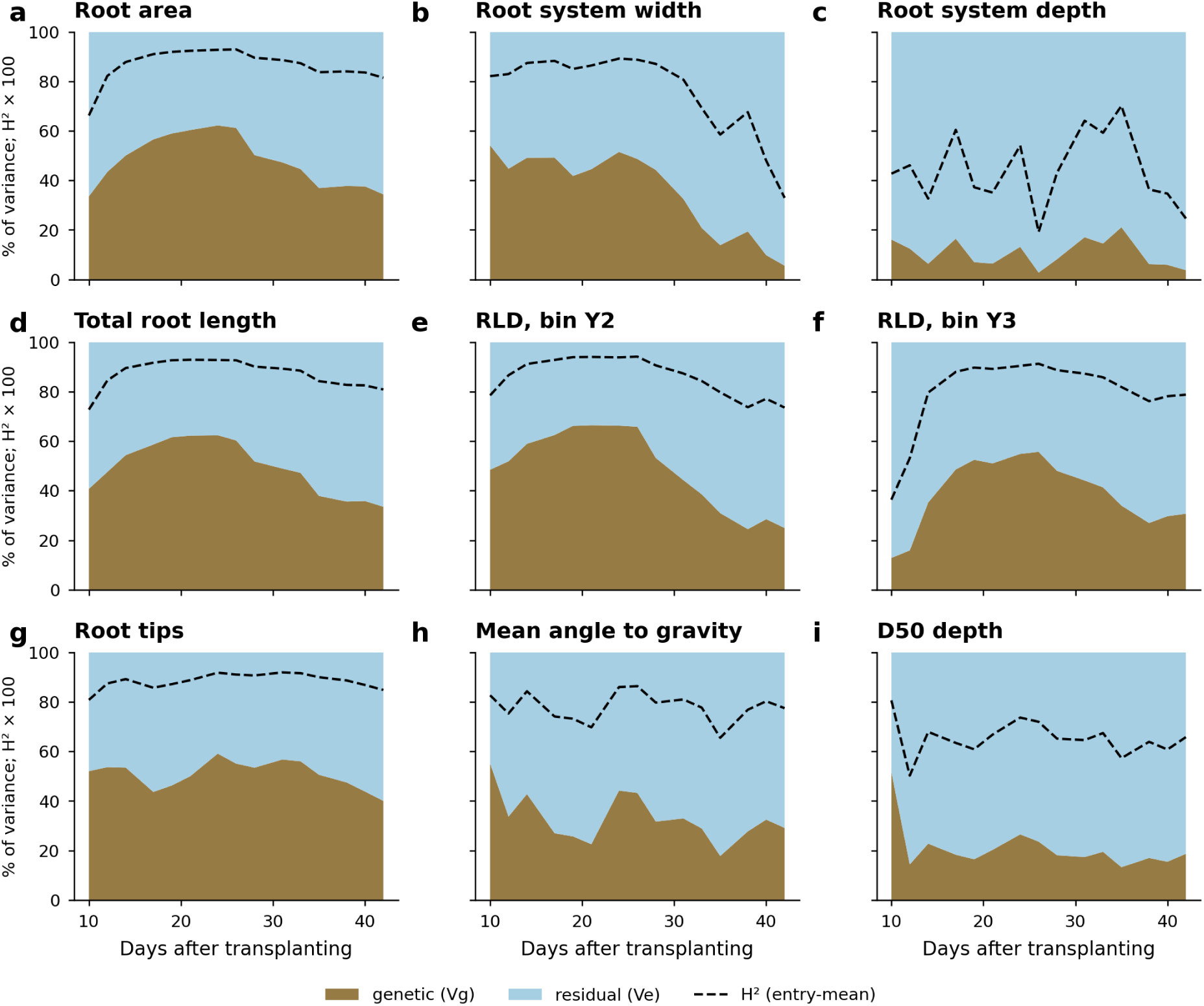
Temporal partitioning of phenotypic variance. Genetic (V_G_) and residual (V_E_) shares of the phenotypic variance, and entry-mean heritability (dashed), estimated at each imaging day for nine traits.

## Supplementary Note S1. Tip-initiated paths and segment support

*Graph.* The skeleton of a root mask is cut at branch points into segments. A segment end is a tip if it is a skeleton node of degree one, as root tips are defined in Trait extraction (this includes the end of the system at the cutting and ends left by gaps in the mask), and a branch point otherwise; segment ends meeting at a branch point form a vertex. A segment is terminal if it has one tip (set E_T_), isolated if it has two, and inner if it is bounded by two branch points (set E_I_). Image coordinates have y increasing downward, so the cutting lies at small y.

*Path.* Every terminal segment e_0_ ∈ E_T_ starts one path P = (e_0_, e_1_,…), entered at its tip; isolated segments start none and are never traversed. Segment directions are taken along the straight line joining a segment’s two ends. At the current vertex u, let d be the unit vector along the last segment, oriented toward u. The candidates are the inner segments f ∈ E_I_ from u to a vertex w that is not lower than u (within a tolerance ε) and not yet on the path; each has a turning angle θ(f) = cos^-1^(d · d_f_), where d_f_ is the unit vector along f from u to w, so that θ = 0° is straight ahead and θ = 180° a reversal. The candidate with the smallest θ is appended (ties by segment index), and the path P ends when no candidate exists or the smallest θ exceeds θ_max_ = 100°. A path is a “route” in the main text.

*Support.* s(e) is the number of paths containing segment e (0 if none). By construction s = 1 for terminal segments, s = 0 for isolated segments, and s ≥ 0 for inner segments; s ≥ 2 marks a segment shared by two or more paths (transport segments in the main text). Figure S3 illustrates the construction.

### Pseudocode

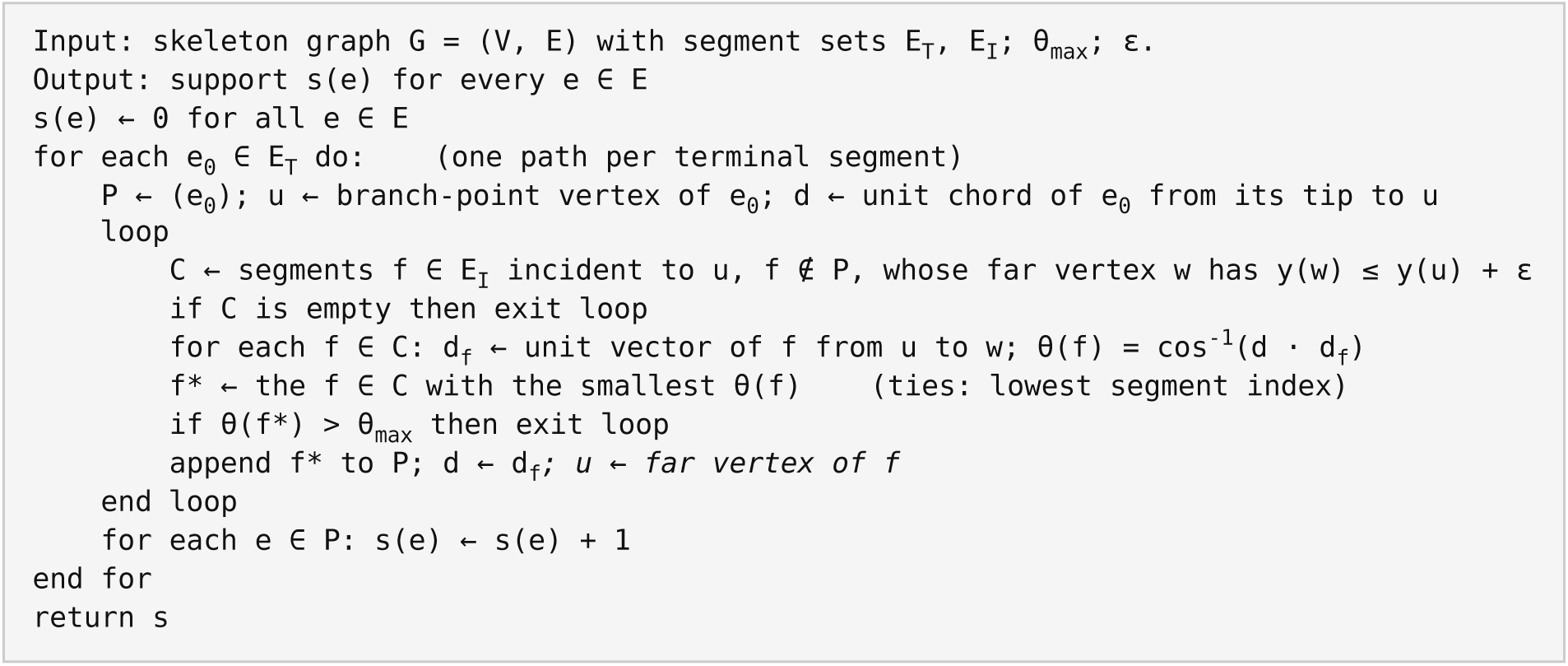

Because every iteration appends a segment not yet in P, every path terminates. The pixel tolerances used to merge segment ends into vertices and for ε, and the skeletonization settings, are given in the code repository (https://github.com/diazgarcialab/grapevine_root_architecture_Munoz2026).

## Supplementary tables

**Table S1.** Number of analyzed plants per genotype and treatment.

**Table S2.** Likelihood-ratio tests from linear mixed models fitted to each image-derived trait (n = 1,108 images from 80 plants). Fixed effects included genotype, imaging day (factor), their interaction, and treatment; plant was treated as a random intercept. Size and count traits were log-transformed. P values are FDR-adjusted across the 21 traits.

**Table S3.** Support-0 fraction and transport-root fraction (over total and over routed length) by genotype.

**Table S4.** Change in root traits between 28 and 38 DAT (the water-deficit period) for 39 well-watered and 37 deficit plants imaged at both dates, and the adjusted treatment difference from a linear model including genotype, the trait value at 28 DAT, and log total root length at 28 DAT. Size traits are changes in log values. RLD gain is the change in root length density in each depth bin (cm cm^-2^). P (FDR) is adjusted across the 23 rows; P (G×T) is the P value of the genotype × treatment interaction.

**Table S5.** Treatment differences in 28→38 DAT changes in the size-overlap subset (53 plants with total root length at 28 DAT within the range shared by both treatments).

**Table S6.** Cell-level colonization models (logistic generalized estimating equations).

**Table S7.** Segment brightness models with and without segment width as a covariate, and restricted to thin segments.

**Table S8.** Standardized associations between root traits and stomatal conductance (313 measurement-image pairs from 80 plants). β is the change in gsw (mol m^-2^ s^-1^) per standard deviation of the trait from a mixed model with treatment, genotype, and date as fixed effects and plant as a random effect. P (FDR) is adjusted across the 22 traits.

**Table S9.** Heritability estimates with bootstrap confidence intervals by trait and imaging day.

**Table S10.** Scan-to-scan variation of traits between consecutive images.

All supplementary tables (S1-S10) are provided in the file Supplementary_Tables.xlsx.

## References

1. van Leeuwen C, Darriet P. The impact of climate change on viticulture and wine quality. J Wine Econ. 2016;11(1):150–167.

2. Gambetta GA, Herrera JC, Dayer S, Feng Q, Hochberg U, Castellarin SD. The physiology of drought stress in grapevine: towards an integrative definition of drought tolerance. J Exp Bot. 2020;71(16):4658–4676.

3. Ollat N, Peccoux A, Papura D, et al. Rootstocks as a component of adaptation to environment. In: Gerós H, Chaves MM, Medrano Gil H, Delrot S, editors. Grapevine in a changing environment: A molecular and ecophysiological perspective. Chichester (UK): Wiley; 2015. p. 68–108.

4. Fort K, Fraga J, Grossi D, Walker MA. Early measures of drought tolerance in four grape rootstocks. J Am Soc Hortic Sci. 2017;142(1):36–46.

5. Fichtl L, Hofmann M, Kahlen K, et al. Towards grapevine root architectural models to adapt viticulture to drought. Front Plant Sci. 2023;14:1162506.

6. Carbonneau A. The early selection of grapevine rootstocks for resistance to drought conditions. Am J Enol Vitic. 1985;36(3):195–198.

7. Chen Y, Fei Y, Howell K, Chen D, Clingeleffer P, Zhang P. Rootstocks for grapevines now and into the future: selection of rootstocks based on drought tolerance, soil nutrient availability, and soil pH. Aust J Grape Wine Res. 2024;2024:6704238.

8. Lynch J. Root architecture and plant productivity. Plant Physiol. 1995;109(1):7–13.

9. LaRue T, Lindner H, Srinivas A, Exposito-Alonso M, Lobet G, Dinneny JR. Uncovering natural variation in root system architecture and growth dynamics using a robotics-assisted phenomics platform. eLife. 2022;11:e76968.

10. Weihs BJ, Heuschele DJ, Tang Z, York LM, Zhang Z, Xu Z. The state of the art in root system architecture image analysis using artificial intelligence: A review. Plant Phenomics. 2024;6:0178.

11. Heinitz CC, Fort K, Walker MA. Developing drought and salt resistant grape rootstocks. Acta Hortic. 2015;1082:305–312.

12. Yildirim K, Yagci A, Sucu S, Tunc S. Responses of grapevine rootstocks to drought through altered root system architecture and root transcriptomic regulations. Plant Physiol Biochem. 2018;127:256–268.

13. Alsina MM, Smart DR, Bauerle T, et al. Seasonal changes of whole root system conductance by a drought-tolerant grape root system. J Exp Bot. 2011;62(1):99–109.

14. Bauerle TL, Smart DR, Bauerle WL, Stockert C, Eissenstat DM. Root foraging in response to heterogeneous soil moisture in two grapevines that differ in potential growth rate. New Phytol. 2008;179(3):857–866.

15. Gambetta GA, Manuck CM, Drucker ST, et al. The relationship between root hydraulics and scion vigour across Vitis rootstocks: what role do root aquaporins play? J Exp Bot. 2012;63(18):6445–6455.

16. Serra I, Strever A, Myburgh PA, Deloire A. Review: the interaction between rootstocks and cultivars (Vitis vinifera L.) to enhance drought tolerance in grapevine. Aust J Grape Wine Res. 2014;20(1):1–14.

17. Smart DR, Schwass E, Lakso A, Morano L. Grapevine rooting patterns: a comprehensive analysis and a review. Am J Enol Vitic. 2006;57(1):89–104.

18. Reingwirtz I, Uretsky J, Cuneo IF, Knipfer T, Reyes C, Walker MA, et al. Inherent and stress-induced responses of fine root morphology and anatomy in commercial grapevine rootstocks with contrasting drought resistance. Plants. 2021;10(6):1121.

19. Cuneo IF, Barrios-Masias F, Knipfer T, Uretsky J, Reyes C, Lenain P, et al. Differences in grapevine rootstock sensitivity and recovery from drought are linked to fine root cortical lacunae and root tip function. New Phytol. 2021;229(1):272–283.

20. Gambetta GA, Fei J, Rost TL, Knipfer T, Matthews MA, Shackel KA, et al. Water uptake along the length of grapevine fine roots: developmental anatomy, tissue-specific aquaporin expression, and pathways of water transport. Plant Physiol. 2013;163(3):1254–1265.

21. Atkinson JA, Pound MP, Bennett MJ, Wells DM. Uncovering the hidden half of plants using new advances in root phenotyping. Curr Opin Biotechnol. 2019;55:1–8.

22. Bak F, Lyhne-Kjærbye A, Tardif S, Dresbøll DB, Nybroe O, Nicolaisen MH. Deep-rooted plant species recruit distinct bacterial communities in the subsoil. Phytobiomes J. 2022;6(3):236–246.

23. McGrail R, Van Sanford DA, McNear DH. Trait-based root phenotyping as a necessary tool for crop selection and improvement. Agronomy. 2020;10(9):1328.

24. Trachsel S, Kaeppler SM, Brown KM, Lynch JP. Shovelomics: high throughput phenotyping of maize (Zea mays L.) root architecture in the field. Plant Soil. 2011;341:75–87.

25. Metzner R, Eggert A, van Dusschoten D, Pflugfelder D, Gerth S, Gerlach N, et al. Direct comparison of MRI and X-ray CT technologies for 3D imaging of root systems in soil: potential and challenges for root trait quantification. Plant Methods. 2015;11:17.

26. Mooney SJ, Pridmore TP, Helliwell J, Bennett MJ. Developing X-ray computed tomography to non-invasively image 3-D root systems architecture in soil. Plant Soil. 2012;352:1–22.

27. Hou L, Gao W, van der Bom F, Weng Z, Doolette CL, Maksimenko A, et al. Use of X-ray tomography for examining root architecture in soils. Geoderma. 2022;405:115405.

28. Teramoto S, Takayasu S, Kitomi Y, Arai-Sanoh Y, Tanabata T, Uga Y. High-throughput three-dimensional visualization of root system architecture of rice using X-ray computed tomography. Plant Methods. 2020;16:66.

29. Dumont C, Cochetel N, Lauvergeat V, Cookson SJ, Ollat N, Vivin P. Screening root morphology in grafted grapevine using 2D digital images from rhizotrons. Acta Hortic. 2016;1136:213–220.

30. Enyew M, Geleta M, Feyissa T, et al. Sorghum genotypes grown in simple rhizotrons display wide variation in root system architecture traits. Plant Soil. 2024;502:87–102.

31. Landl M, Schnepf A, Vanderborght J, et al. Measuring root system traits of wheat in 2D images to parameterize 3D root architecture models. Plant Soil. 2018;425:457–477.

32. Rellán-Álvarez R, Lobet G, Lindner H, et al. GLO-Roots: an imaging platform enabling multidimensional characterization of soil-grown root systems. eLife. 2015;4:e07597.

33. Pornaro C, Macolino S, Menegon A, Richardson M. WinRHIZO technology for measuring morphological traits of bermudagrass stolons. Agron J. 2017;109(6):3007–3010.

34. Yasrab R, Atkinson JA, Wells DM, French AP, Pridmore TP, Pound MP. RootNav 2.0: deep learning for automatic navigation of complex plant root architectures. GigaScience. 2019;8(11):giz123.

35. Liu J, An F, Yuan K, Chen QB, Wang ZH. Application of SmartRoot system for determining morphological parameters of fine roots of Hevea brasiliensis. Chin J Plant Ecol. 2013;37(8):786–792.

36. Seethepalli A, Dhakal K, Griffiths M, Guo H, Freschet GT, York LM. RhizoVision Explorer: open-source software for root image analysis and measurement standardization. AoB Plants. 2021;13(6):plab056.

37. Zheng P, Gao D, Fan DP, et al. Bilateral reference for high-resolution dichotomous image segmentation. CAAI Artif Intell Res. 2024;3:9150038.

38. van der Walt S, Schönberger JL, Nunez-Iglesias J, et al. scikit-image: image processing in Python. PeerJ. 2014;2:e453.

39. Bradski G. The OpenCV library. Dr Dobb’s J Softw Tools. 2000;25(11):120–125.

40. Nunez-Iglesias J, Blanch AJ, Looker O, Dixon MW, Tilley L. A new Python library to analyse skeleton images confirms malaria parasite remodelling of the red blood cell membrane skeleton. PeerJ. 2018;6:e4312.

41. Seabold S, Perktold J. statsmodels: Econometric and statistical modeling with Python. In: Proceedings of the 9th Python in Science Conference (SciPy 2010); 2010. p. 92–96.

42. Smith AG, Lamprinidis S, Seethepalli A, et al. A systematic comparison of transformers and ConvNets for root segmentation across nine datasets. Plant Methods. 2026;22:52.

43. Fichtl L, Leitner D, Schnepf A, Schmidt D, Kahlen K, Friedel M. A field-to-parameter pipeline for analyzing and simulating root system architecture of woody perennials: Application to grapevine rootstocks. Plant Phenomics. 2024;6:0280.

44. Mahmud KP, Field SK, Rogiers SY, Nielsen S, Guisard Y, Holzapfel BP. Rootstocks alter the seasonal dynamics and vertical distribution of new root growth of Vitis vinifera cv. Shiraz grapevines. Agronomy. 2023;13(9):2355.

45. Zobel RW, Waisel Y. A plant root system architectural taxonomy: a framework for root nomenclature. Plant Biosyst. 2010;144(2):507–512.

46. Freschet GT, Pagès L, Iversen CM, Comas LH, Rewald B, Roumet C, et al. A starting guide to root ecology: strengthening ecological concepts and standardising root classification, sampling, processing and trait measurements. New Phytol. 2021;232(3):973–1122.

47. Alonso-Forn D, Buesa I, Flor L, Sabater A, Medrano H, Escalona JM. Implications of root morphology and anatomy for water deficit tolerance and recovery of grapevine rootstocks. Front Plant Sci. 2025;16:1541523.

48. Barrios-Masias FH, Knipfer T, McElrone AJ. Differential responses of grapevine rootstocks to water stress are associated with adjustments in fine root hydraulic physiology and suberization. J Exp Bot. 2015;66(19):6069–6078.

49. Vargas S, Osorio N, González A, Ortega-Farías S. Effect of irrigation management on the relationship between stomatal conductance and stem water potential on cv. Cabernet Sauvignon. BIO Web Conf. 2023;56:01012.

50. Tandonnet JP, Marguerit E, Cookson SJ, Ollat N. Genetic architecture of aerial and root traits in field-grown grafted grapevines is largely independent. Theor Appl Genet. 2018;131(4):903–915.

51. Blois L, de Miguel M, Bert PF, et al. Dissecting the genetic architecture of root-related traits in a grafted wild Vitis berlandieri population for grapevine rootstock breeding. Theor Appl Genet. 2023;136(11):223.

52. Dudley SE, Diaz-Garcia L, Uyehara IK, McElrone AJ, Cannizzaro A, Sharma S, et al. Grapevine adventitious root traits vary based on propagation method. OENO One. 2026;60(2):9599.

53. Bernardo S, Marguerit E, Ollat N, Gambetta GA, de Saint Cast C, de Miguel M. Root system ideotypes: what is the potential for breeding drought-tolerant grapevine rootstocks? J Exp Bot. 2025;76(11):2970–2984.

